# Improved detection of microbiome-disease associations via population structure-aware generalized linear mixed effects models (microSLAM)

**DOI:** 10.1101/2024.06.27.600934

**Authors:** Miriam Goldman, Chunyu Zhao, Katherine S. Pollard

## Abstract

Microbiome association studies typically link host disease or other traits to summary statistics measured in metagenomics data, such as diversity or taxonomic composition. But identifying disease-associated species based on their relative abundance does not provide insight into why these microbes act as disease markers, and it overlooks cases where disease risk is related to specific strains with unique biological functions. To bridge this knowledge gap, we developed microSLAM, a mixed-effects model and an R package that performs association tests that connect host traits to the presence/absence of genes within each microbiome species, while accounting for strain genetic relatedness across hosts. Traits can be quantitative or binary (such as case/control). MicroSLAM is fit in three steps for each species. The first step estimates population structure across hosts. Step two calculates the association between population structure and the trait, enabling detection of species for which a subset of related strains confer risk. To identify specific genes whose presence/absence across diverse strains is associated with the trait, step three models the trait as a function of gene occurrence plus random effects estimated from step two. Applying microSLAM to 710 gut metagenomes from inflammatory bowel disease (IBD) samples, we discovered 49 species whose population structure correlates with IBD. In addition, after controlling for population structure, we found 57 microbial genes that are significantly more common in healthy individuals and 26 that are more common in IBD patients, including a seven-gene operon in *Faecalibacterium prausnitzii* that is involved in utilization of fructoselysine from the gut environment. Overall, microSLAM detected IBD associations for 45 species that were not detected using relative abundance tests, and it identified specific strains and genes underlying IBD associations for 13 other species. These findings highlight the importance of accounting for within-species genetic variation in microbiome studies.

**Author Summary:** The species composition of the human gut microbiome differs significantly between individuals and is associated with various diseases. Many studies have sought to understand this relationship by examining the relative amount of each bacterial species within metagenomic sequencing data from sick and healthy individuals. However, this approach makes it challenging to pinpoint which genes and pathways of a disease-associated species might actually contribute to disease risk, and it misses species where only certain strains are disease associated. To overcome these challenges, we developed an R package, called microSLAM, that uses mixed-effects modeling to associate within-species genetic variation to host traits. microSLAM performs two types of tests for each species: one for identifying strain-disease associations and another for identifying gene-disease associations. The gene tests account for the genetic relatedness of strains across hosts, making them particularly useful for detecting mobile genes. We applied microSLAM to hundreds of gut metagenomes from inflammatory bowel disease studies, identifying dozens of novel associations that were missed using relative abundance tests. MicroSLAM is a general modeling approach that can be applied to human traits beyond disease case/control studies and to microbiomes from other environments.

## Introduction

The human body is home to a complex community of microorganisms, known as the microbiome, which encodes millions of genes [1]. The species composition of the microbiome differs significantly between individuals and is associated with host genetics, diet, immune system, and several human diseases [2–5]. As microbiome species evolve, individual lineages lose and gain genes through horizontal gene transfer [6,7] and other processes that create structural variation [8–10]. The resulting pangenome can be quantified from shotgun metagenomics data [11–13], which has revealed immense genetic diversity between and within human hosts [14]. Even when two people harbor the same microbial species, the cells within those populations are likely to perform different functions [10,15]. For example, prior studies identified many cases of variable virulence and antibiotic resistance [16,17], a set of pro-inflammatory genes from specific strains of *Ruminococcus gnavus* [18], a *Faecalibacterium prausnitzii* GalNAc utilization pathway linked to with cardiometabolic health [19], and a strain of *Escherichia coli* with enhanced ability to live on the intestinal mucus that is associated with inflammatory bowel disease (IBD) [20]. These findings underscore the limitations of using species abundance alone to gain insight into host-microbiome interactions.

In this study, we consider two ways to leverage within-species pangenomic diversity to discover associations between the microbiome and a trait of the host, such as disease. The first is designed for when a species has a strain or group of related strains that predicts the trait. Identifying and isolating trait-associated strains facilitates experimental investigations into host-microbiome interactions, and strains enriched in healthy hosts have been proposed as components of probiotics and therapies [21–24]. Due to the systematic structure of bacterial genomes in which many genes have correlated presence/absence across strains–especially closely related strain–this approach will typically identify a large set of trait-associated genes.

While any of these genes could be a good biomarker (e.g., for diagnosis or patient stratification), most of them are not good candidates for follow-up studies of causal mechanisms. Therefore, we also consider a second case in which one or a small number of individual genes predict the trait. Such associations are easiest to detect if the genes are rapidly gained and lost (e.g., mobile elements), so that they associate with the trait independently of evolutionary relationships amongst strains. Genes like this are promising candidates for discovering causal mechanisms through which microbes modify host health and treatment responses.

To identify trait-associated microbiome strains and genes, we developed a statistical model that can be used to perform a metagenome-wide association study (MWAS) for any continuous or binary host trait. Building on the work done on generalized linear mixed-effects models from human genetics [25–28], this modeling approach uses gene presence/absence data from cohorts with metagenomic sequencing to first estimate a between-sample genetic relatedness matrix (GRM) for each microbiome species and associate this population structure with the host trait. Then, each gene in a species’ pangenome is tested for its trait association after accounting for the relatedness of strains across hosts using random effects derived from the GRM. Our methodology is implemented in an open-source R package, called microSLAM, which can be used with quantitative and binary traits (including unbalanced case/control studies), scales to thousands of samples, and has a controlled type one error rate. The two tests in microSLAM enable researchers to detect new associations and to refine associations discovered using relative abundance.

To investigate the utility of microSLAM, we analyzed a compendium of 710 publicly available metagenomes from IBD case/control studies (S1 Table). IBD is an inflammatory condition of the gastrointestinal tract characterized by its persistence [29]. IBD afflicts roughly three million Americans [30], and its incidence has continued to increase in older adults in recent years [31]. The gut microbiome has long-standing links to IBD, including species abundance and gene associations [15,20,29,32–41]. Here, we combined MIDAS v3 pangenome profiling [11] with microSLAM to quantify associations between IBD and relative abundance, population structure, and gene presence/absence across 71 common members of the human gut microbiome. These analyses identified 49 species with IBD-associated population structure and 83 significant gene families, which we interpreted at the pathway level within and across microbiome species. Tests based on relative abundance would have missed these associations.

## Results

### MicroSLAM modeling approach

We present a new method, called microbiome population Structure Leveraged Association Model (microSLAM). The examples in this study focus on the human gut microbiome and IBD case/control cohorts, but microSLAM can be applied to any trait or environment. MicroSLAM implements a generalized linear mixed-effects model that enables two complementary statistical tests of association between a host trait and within-species genetic variation (Fig 1A; Methods). Both tests use the presence/absence of genes from a given species’ pangenome across samples, which can be quantified from metagenomic sequencing data using tools such as MIDAS v3, panX, and Roary [11–13]. The inputs are a gene presence/absence matrix, any covariates one wishes to include, and the trait data, which can be quantitative or binary (e.g., case/control). The microSLAM method is implemented as an open-source R package at: https://github.com/miriam-goldman/microSLAM.

**Figure 1.**
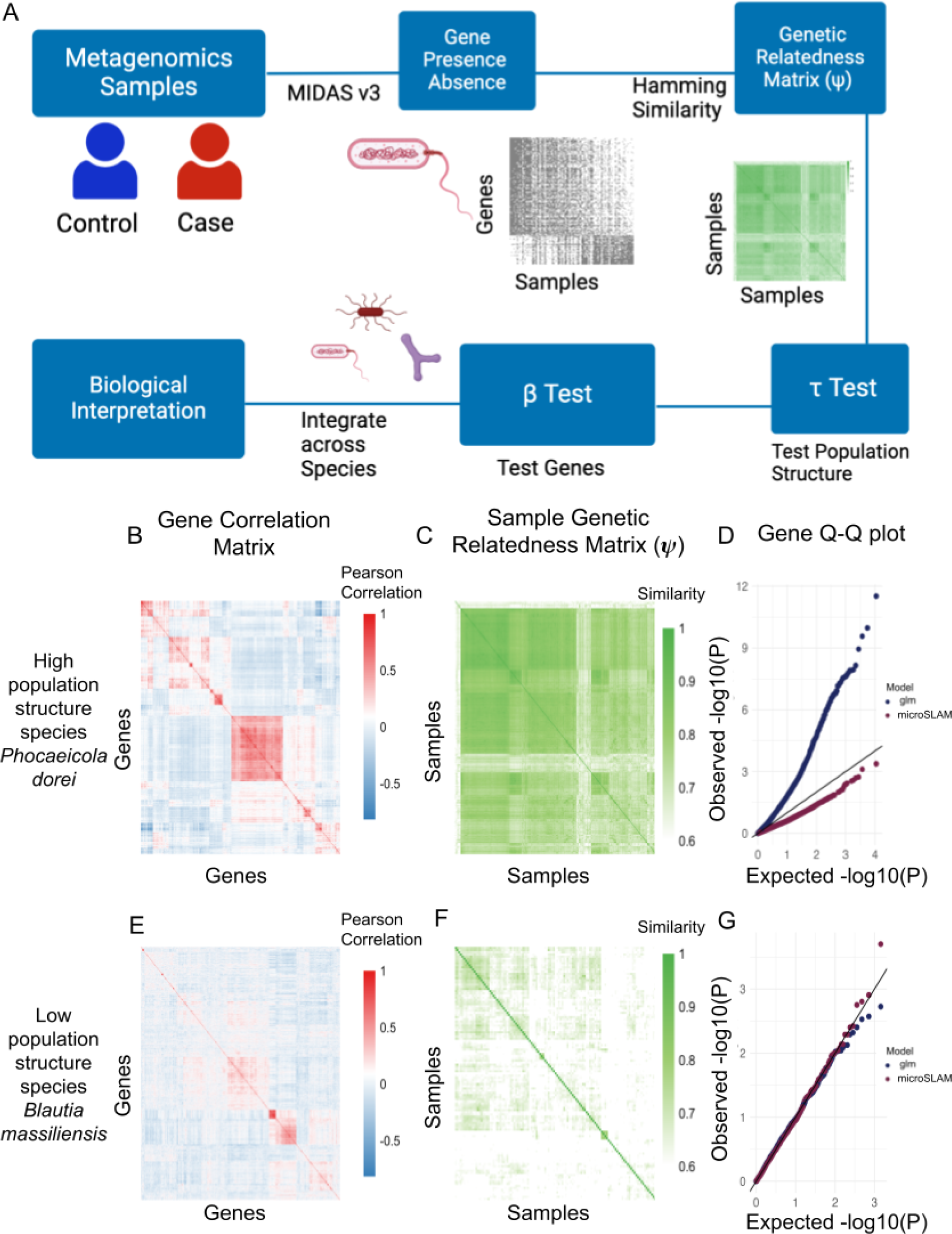
MicroSLAM motivation and approach. A) Flow chart of microSLAM modeling approach (diagram created in BioRender). B – G) Two bacterial species with different population structures. First row: *Phocaeicola dorei* (260 IBD cases, 218 controls); Second row: *Blautia massiliensis* (73 IBD cases, 44 controls). B&E) Heatmap of gene by gene correlation matrix based on gene presence/absence across IBD samples. Red: high positive correlation, Blue: high negative correlation. C&F) Heatmap of sample by sample genetic relatedness matrix (1 minus Hamming distance of gene presence/absence profiles). Dark green: high similarity, White: low similarity. D&G) Q-Q plot for p-values from tests of association between case/control status and presence/absence of individual genes in the pangenome. Tests are based on micoSLAM and standard logistic regression that does not adjust for population structure (glm). The diagonal line shows expected p-values under the null hypothesis of no association. Pangenome profiling for the metagenomes was done using MIDAS v3.

The first test, the τ test, identifies species for which trait variation is associated with variation in overall gene content, as quantified by a random effect (*b_i_*) for each host that is estimated using a GRM (1-Hamming distance of gene presence/absence) and quantifies the association between the sample’s lineage and the trait. We refer to the output of the τ test as strain-level associations, because many gene families which are jointly present/absent across hosts are all equally likely to be associated with the trait. Identifying strain-trait associations is important because this improves the precision of research and therapeutics development based on cultured strains beyond simply picking a random strain from a trait-associated species, which may in fact not be one of the strains driving the species-level association. The genes jointly defining trait-associated strains (i.e., those positively or negatively associated with *b*) also provide signatures that can be used for predictive modeling and potential diagnostics.

However, the many jointly evolving genes that define trait-associated strains will make it difficult to pinpoint causal mechanisms; some of them may play direct roles in the etiology of the trait or in enabling the strain to survive in hosts with that trait, while others are simply present in the same lineage. To address this, microSLAM implements a second test, called the β test, that identifies individual gene families which are significantly associated with the trait above and beyond what is expected given the GRM. This is accomplished by modeling the trait as a function of each gene family’s presence/absence with a generalized linear mixed effects model that includes the random effect (*b_i_*) for each sample (Methods). The resulting significant gene families may be recently and/or recurrently lost and gained (e.g., via mobile elements). To be detected, they must be evolving somewhat independently of the gene families that distinguish strains and in patterns that strongly associate with the host trait. These are high-confidence candidates for studying causal mechanisms. Going beyond standard species relative abundance tests, microSLAM’s two within-species tests are designed to enable (i) identification of specific strains and gene functions driving species-trait associations, and (ii) detection of novel trait associations not detectable at the species level.

### Population structure in inflammatory bowel disease gut microbiomes

We compiled 710 publicly available gut metagenomes from five inflammatory bowel disease (IBD) case/control studies (S1 Table) and performed pangenome profiling of them using MIDAS v3. There were 71 species with sufficient sequencing coverage to analyze within species genetic variation. After dropping gene families that are nearly always present or nearly always absent (Methods), we had an average of 2,254 gene families per species and a total of ∼160 thousand across species. All species showed some population structure. In some species, such as *Phocaeicola dorei (P. dorei),* many gene families are co-evolving and show a high correlation in their presence/absence across hosts (Fig 1B). In turn, we see two distinct subgroups of strains in the GRM (Fig 1C). This high level of structure might be the result of selection pressures, drift, or a recent population expansion. When we perform MWAS for all *P. dorei* gene families using logistic regression (glm), we observe that most genes are significantly associated with IBD case/control status (Fig 1D). This inflation is similar to the well-known problem in human genetics in which ancestry-associated variants are all highly significant when genetic ancestry differs between cases and controls [42]. In contrast, the gene-level test in microSLAM does not show inflation, because our model adjusts for population structure when testing individual gene families for disease associations. We therefore hypothesize that inflation is a consequence of high population structure resulting from a high correlation between gene families. Supporting this, *Blautia massilensis* does not have many genes that are correlated (Fig 1E) and shows less structure in its GRM (Fig 1F). Accordingly, the glm p-values do not show inflation, and the microSLAM output is very similar to that of glm.

These results suggest that if we wish to identify individual gene families with unexpectedly high associations with a host trait given the species’ GRM, the mixed modeling approach in microSLAM provides a way to adjust for population structure across hosts, just as mixed models have enabled human geneticists to account for confounding from genetic ancestry. However, population structure is not necessarily a confounder in microbiome research, and it may also be of interest to identify trait-associated strains, defined by the presence and absence of many gene families relative to other strains. These genes would not be significant in the microSLAM β test, because they are highly correlated with population structure. For this reason, microSLAM also includes a strain-level test, the τ test.

Consider, for example, *Ruminococcus B gnavus (R. gnavus),* a species that has long been associated with IBD [18,43,44]. The *R. gnavus* GRM shows two distinct groups when hosts are sorted based on their *b*_*i*_ values (Fig 2A). One of these groups only contains IBD individuals, while the other is split between IBD and controls. The *b_i_* estimates better separate cases and controls than do the first two principal coordinates of the gene presence/absence matrix (Fig 2B). Not surprisingly, when we apply the microSLAM τ test to *R. gnavus*, we obtain a large and statistically significant measure of association (τ=4.67, permutation p-value=0.0001; Fig 2C), and the resulting model can classify IBD cases with high accuracy (ROC AUC = 0.987; Fig 2D). Now, if we look at the genes that are most highly correlated to the estimated *b_i_* values of the samples, we identify 238 out of the ∼1200 non-core, non-rare genes used in the analysis that are all nearly equally associated with IBD (Fig 2E). None of these genes are significantly associated with IBD after adjusting for population structure. Thus, we were able to identify several hundred highly correlated genes that form a predictive signature for the *R. gnavus* strains present in IBD patients versus controls. These observations illustrate the importance of including the τ test in microSLAM.

**Figure 2.**
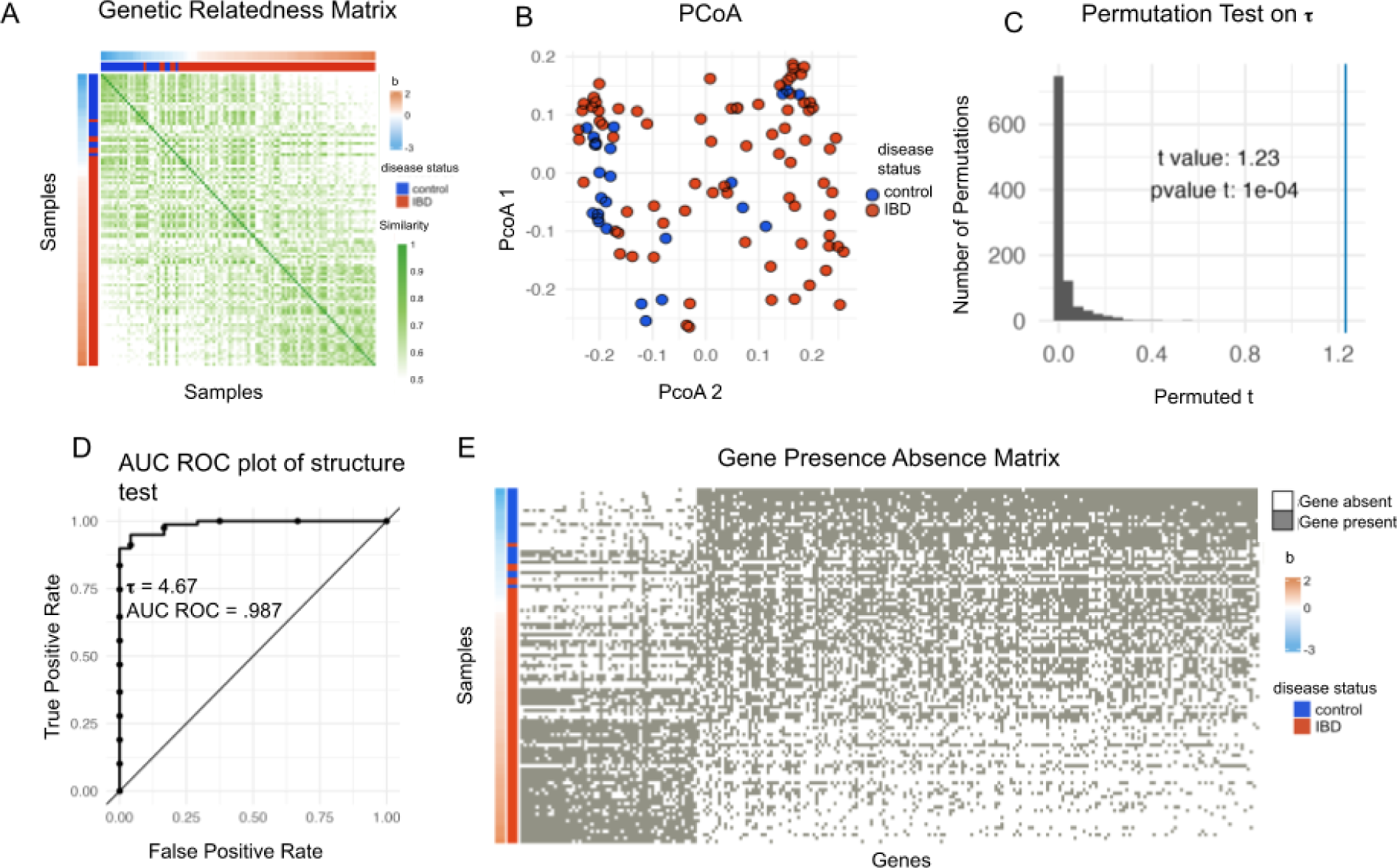
MicroSLAM detects both strain and gene associations. A) The GRM for *Ruminococcus B gnavus* with hosts sorted by their estimated *b* values and annotated by their disease status. B) PCoA from the *R. gnavus* gene presence/absence colored by host disease status (as in A). C) Histogram of permutation test statistics (t-values) from the τ test for *R. gnavus.* The line denotes the observed value of t. D) ROC plot for the microSLAM τ test model for *R. gnavus.* The statistic τ quantifies population structure. E) Gene presence/absence plot for a subset of genes associated with the random effect *b* for *R. gnavus.* Samples are ordered by *b* and annotated by their disease status.

### MicroSLAM controls false positive rates and increases specificity in simulations

The examples in Figs 1 and 2 suggest that microSLAM’s τ test can detect strain-trait associations when species have a high degree of population structure and its β test may control false positive gene-disease associations better than a standard glm, albeit somewhat conservatively. But the ground truth is unknown in real data. Hence, we designed a series of simulations to assess the performance of both of microSLAM’s tests. Our simulation strategy leveraged the IBD compendium in order to capture the range of patterns observed in real data, while varying parameters such as effect size and sample size. For the β test, we compared microSLAM to glm in order to evaluate the effects of adjusting for population structure via the random effects *b_i_*.

First, we evaluated the τ test. To the best of our knowledge, there is no other method that performs this type of association test, so we did not compare microSLAM to alternative approaches. To quantify type 1 error (i.e., false positive rate), we simulated gene presence/absence matrices with core and accessory genes, as well as a set of strain-specific genes, and for each host a binary trait was simulated independently of the gene presence/absence matrix so that the GRM is not associated with the trait (τ test simulation simulation 1; Methods). We investigated a sample size of 100 hosts, which is on the low end of what we observed for species in the IBD compendium, and repeated the simulation 1000 times, keeping track of how many iterations had a permutation p-value < 0.05. We observed a false positive rate of 0.054, which is very close to the expected value of 0.05. This indicates that the false positive rate of microSLAM’s τ test is approximately correct.

To evaluate the power of the τ test, we modified the prior simulation so that the trait depends on presence/absence of a particular strain (τ test simulation 2; Methods). We varied the strength of the strain-trait association (odds ratio) and explored sample sizes ranging from 60 to 250. As expected, power increases with the odds ratio and sample size (Fig 3B). MicroSLAM achieves ∼80% power at an odds ratio of 1.5 with 250 samples, whereas an odds ratio greater than 2.0 is needed for similar power with only 60 samples. These results provide practical guidelines for the expected performance of the τ test.

**Figure 3.**
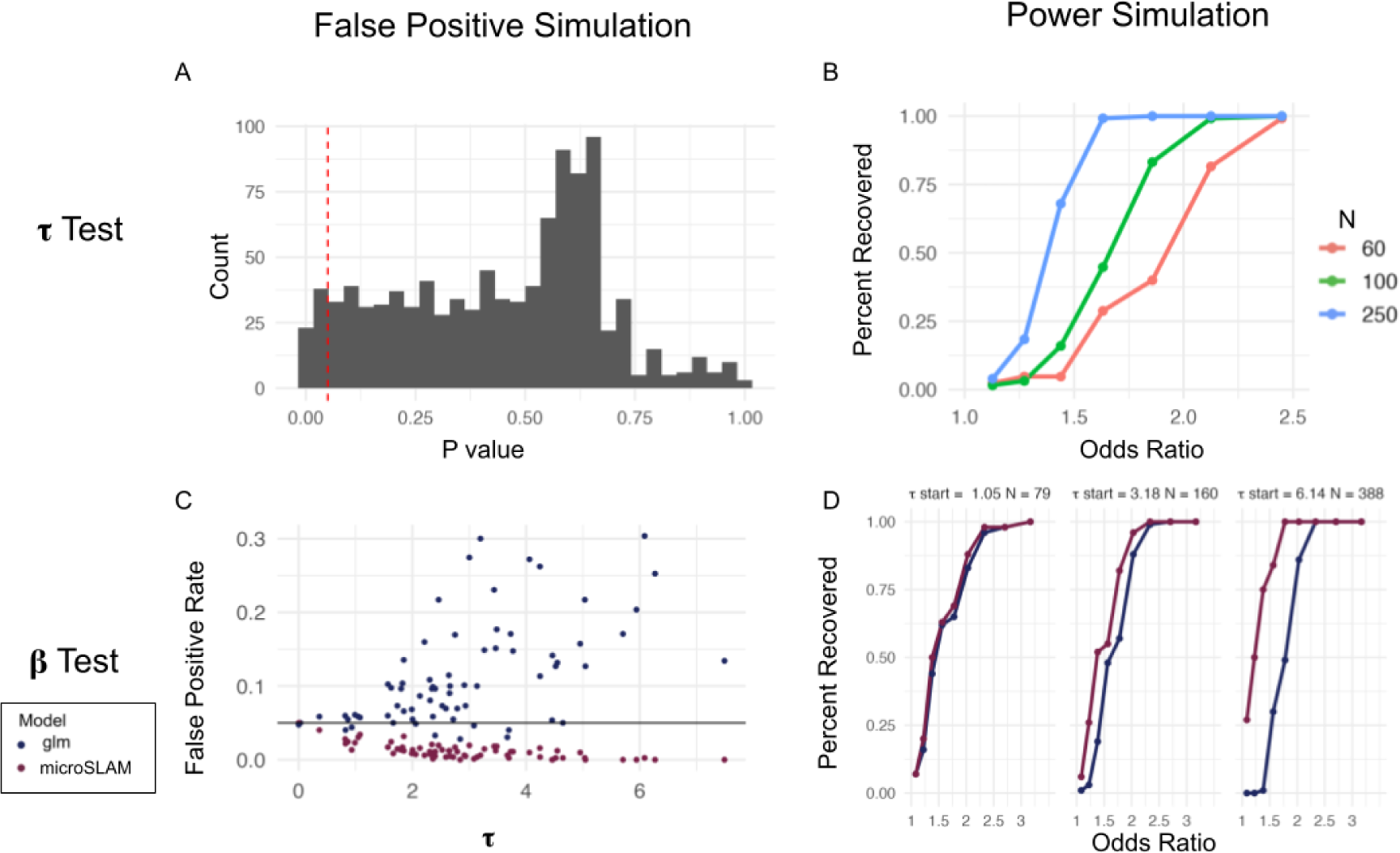
Simulations show that microSLAM improves power and false positive rates. A) The false positive rates of the τ test of microSLAM were estimated using simulations with varying GRMs but no trait associations. We simulated gene presence/absence and GRMs for the 1000 iterations (τ test simulation 1; Methods). A histogram of p-values for the τ tests shows that the percentage of tests with a p-value < 0.05 is 5.4%. B) Power of the τ test for simulations with a range of values for the odds ratio of the simulated y compared to presence of the trait-associated strain (τ test simulation 2; Methods), repeated for different numbers of samples (N). C) False positive rates of the β tests for glm and microSLAM were estimated using simulations with varying levels of population structure (τ) but no trait associations. We simulated gene presence/absence using the GRMs for the 71 species in the IBD compendium (β test simulation 1; Methods). The false positive rate increases with τ for the glm and is generally above the targeted level (0.05; horizontal line), while it decreases and is generally below 0.05 for microSLAM. D) Power for 3 simulated species with different τ values and numbers of samples (N). For a subset of genes, presence/absence is simulated based on the trait using a range of odds ratios; other genes have presence probabilities that do not depend on the trait (β test simulation 2; Methods).

Next, we investigated the type 1 error and power of microSLAM’s β test compared to glm, which is the standard approach used in the literature. We considered the case where gene presence/absence is not associated with a binary trait, but it is associated with population structure, and hence genes are correlated with each other. To do so, we simulated gene presence/absence using principal components of the observed GRM for each of the 71 species in the IBD compendium (β test simulation simulation 1; Methods). MicroSLAM controlled the false positive rate below 0.05 for all but two species where it is exactly 0.05 (*Dorea A longicatena, Roseburia sp900552665*). In contrast, the glm without a random effect adjusting for population structure failed to do so for all but 9 species (*Faecalibacterium prausnitzii, Bifidobacterium adolescentis, Bariatricus comes, Blautia A faecis, Faecalibacillus intestinalis, Gemmiger qucibialis, Akkermansia muciniphila, Roseburia sp900552665, Acetatifactor sp900066565*), a failure rate of 87.3% (62/71). In addition, as the estimated τ increased, the false positive rate of the glm dramatically increased while the false positive rate for microSLAM decreased slightly (Fig 3C).

To explore if this conservative control of the false positive rate affects the power of microSLAM’s β test, we performed simulations where 100 true positive genes are added to the previously stimulated genes, meaning that they have a presence/absence pattern that is associated with the simulated trait (β test simulation 2; Methods). We varied the strength of the association (odds ratio) and evaluated power at an empirical false positive rate of 0.05 (calculated using the non-trait-associated genes). These analyses show that microSLAM consistently has either the same or higher power than the glm at the same false positive rate (Fig 3D), with the difference between methods being most pronounced with a higher number of samples and a high degree of population structure (τ) (Fig S1). In order to understand exactly what types of genes lead to the higher false positive rate we simulated a data set completely *de novo* without using the IBD compendium (β test simulation 3; Supplement). This showed that the strain-associated genes tended to lead to an increase in false positives for the glm, while microSLAM was able to differentiate the true positives from genes linked to the strain but not directly associated with the trait (Fig S2). Thus microSLAM’s β test has better specificity than does a standard glm.

### MicroSLAM reveals IBD associations across 71 gut microbiome species

We next sought to examine associations in our IBD compendium using microSLAM’s population structure and gene tests. First, we performed the standard species-level analysis in which the relative abundance of each of the 71 gut species (quantified using kraken2 and bracken [45–47]; Methods) is tested for association with IBD case/control status using logistic regression, adjusting for host age. We also explored adjusting for study, but found that no species had significant study effects. Medications and other clinical covariates are important confounders but unfortunately were not provided in the publicly available datasets. We found that 13/71 species (18%) had significant relative abundance associations (localFDR<10%; Fig 4A).

**Figure 4.**
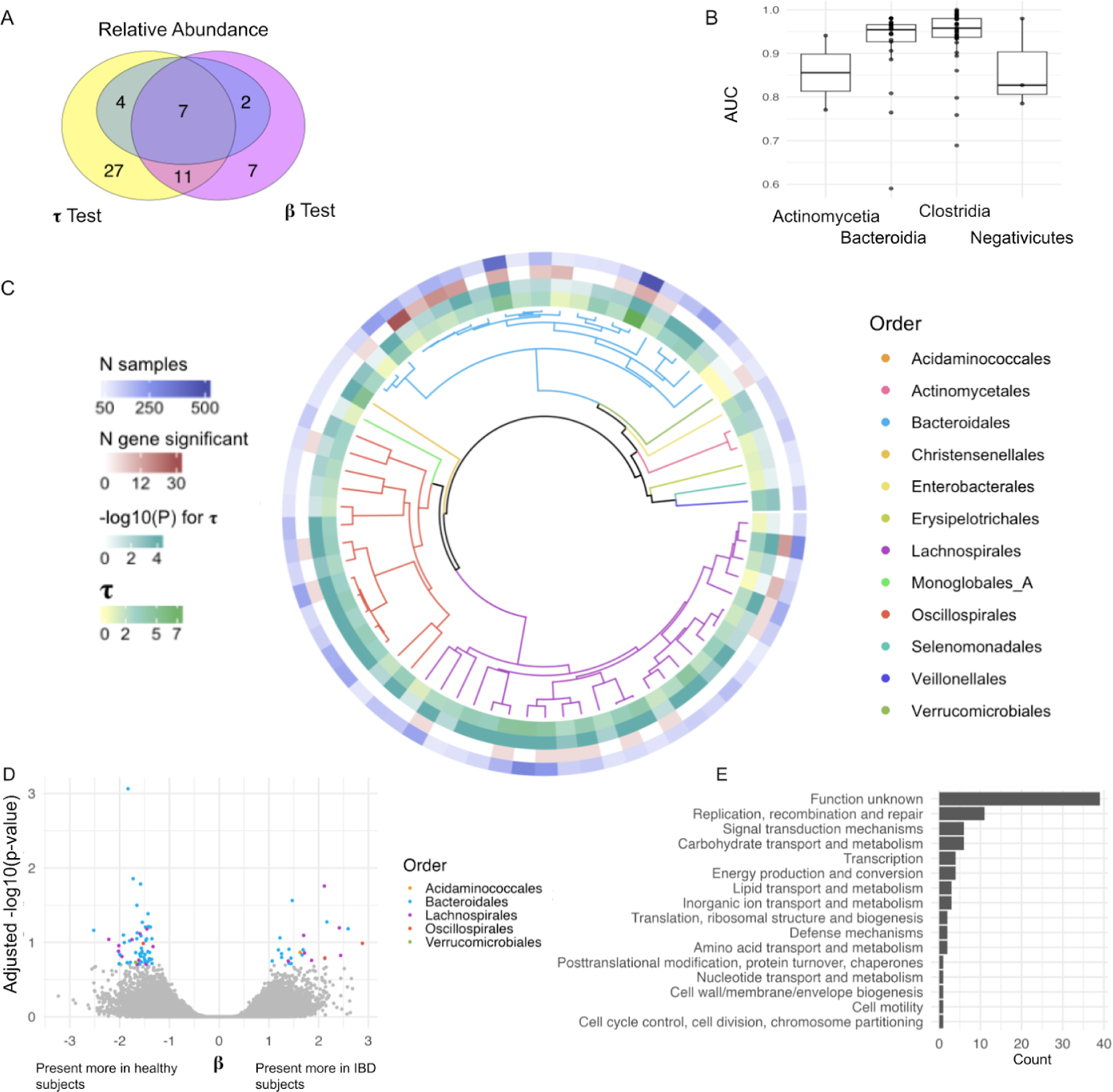
MicroSLAM identifies novel IBD associations. We analyzed all 71 species in our IBD compendium for three types of associations with case/control status: relative abundance (kraken2+braken: amount of the species predicts disease), population structure (microSLAM τ: strain predicts disease), and gene family (microSLAM β: gene presence/absence predicts disease). A) Venn diagram showing the number of species with significant IBD associations of each type. For genes, we counted the species if it had at least one significant gene family; species varied in the number of hits (S2 Table). All tests are localFDR adjusted for multiple testing. B) Boxplots showing the AUC ROC from τ test models for all 71 species, stratified by bacterial class. C) UHGG species tree for all 71 species, colored by order. The τ value, p-value for τ test, number of significant genes, and number of samples for each species are plotted in the outer rings. D) Volcano plot for β tests with significant genes (localFDR < 0.2) colored by bacterial order. (S3 Table) E) Bar plot of COG categories for the 83 genes with significant β tests.

To investigate strains potentially responsible for these relative abundance associations, and to explore the possibility that some species have strain associations without relative abundance associations, we next ran microSLAM using each species’ gene presence/absence matrix and corresponding GRM (quantified using MIDASv3 [11]; Methods). We used microSLAM’s τ test to identify species whose population structure is associated with IBD case/control status. At localFDR<10%, 49/71 species (69%) were significant, meaning that cases and controls tend to harbor distinct strains consistently across studies. Thirty-four of these species were not detected in the relative abundance test (Fig 4A). Twenty-seven of the species were only detected from the population structure test, of those 18/27 (67%) are from class *Clostridia*. These are well-powered studies, each including both cases and controls, and they come from different geographical regions with different diets and lifestyles. Furthermore, the metagenomics data was generated using DNA library preparation methods, which can introduce batch effects. Hence, detecting strain-disease associations across studies suggests that these associations are truly linked to microbial population structure, rather than an unmeasured confounder (S3 Fig), though we cannot rule out confounding due to the limited amount of publicly available data about the study subjects. As opposed to simply including a PC from the GRM to represent the structure of the population, the population structure component of the model pulls out from the GRM the cryptic relatedness that can be attributed to the phenotype given the covariates that are included. If we were to have more information about host covariates (e.g., diet or exercise), then the population structure component would be the portion of the GRM that can be attributed to the phenotype given these covariates of the host.

In addition to assessing the statistical significance of τ via a permutation test, we also report the area under the curve (AUC) for the receiver operating characteristic (ROC) from the τ test model. This shows how well the population structure component is able to separate the cases from the controls. The AUC is calculated within the same training data, because the random effects *b,* which are per-host parameters (i.e., on per subject) generated from the GRM, are unknown for new hosts and hence the fitted model does not generalize beyond the training set. Overall the AUC from the τ tests was quite high; 54 species had AUC over 0.9. Class *Clostridia* tended to have the highest AUC values and a smaller variance in AUC values compared to *Bacteroidia* (Fig 4B). In addition to *R. gnavus* (Fig 2), species with significant τ tests included *Agathobacter rectalis* (previously found to be related to IBD under certain conditions [48]) and *Phocaeicola coprocola* (formally *Bacteroides coprocola*, which has been shown to have a relationship with ulcerative colitis [49]). In both of these species, there were no significant genes with the β test, but with the information from the τ test genes differentiating IBD-associated strains can be identified.

To investigate specific gene families associated with IBD case/control status, above and beyond the genes that define IBD-associated strains, we next applied microSLAM’s β test. Across the 71 species, 83 genes from 27 species showed significant associations after adjusting for population structure (localFDR<20%, which is the threshold with optimal lift and somewhat more lenient than the 10% threshold used for the other two tests). Seven of these species did not have significant relative abundance or population structure associations, underscoring the unique information captured by each of microSLAM’s tests (Fig 4A).

Having analyzed IBD associations at the species, strain and gene level, we integrated these results across the 71 species to look for phylogenetic trends (Fig 4C). Out of the 71 analyzed gut species, 13 (spread across the phyla *Firmicutes A*, *Firmicutes C*, *Bacteriodota*, and *Actinobacteria*) had no significant IBD associations, possibly due to a lower number of samples (N<100 for ten species). Of the 13 species with relative abundance associations, all were detected on one or both of microSLAM’s tests (Fig 4A), suggesting that relative abundance differences are often accompanied by differences in gene content. Looking across the phylogenetic tree, Lactobacillus species tend to have the least IBD-associated population structure (low values of τ), although there is a subclade of two species with higher τ values. On the other hand, Oscillospirales tend to have high values of τ, and most species in this order do not have any significant genes. Finally, Bacteroidales stands out as the order with the most significant genes (60/83), consistent with species in this order having many mobile and accessory genes [50].

To further explore the functions of genes identified by microSLAM’s β test (Fig 4D), we ran multiple gene annotation pipelines. As expected, most of the 83 significant genes had no functional annotation. For example, 39/83 are in the EggNOG COG category “function unknown”. The remaining annotated genes were too few in number to perform well-powered enrichment analyses, but we did note several interesting trends (Fig 4E). The most common COG, encompassing 11 genes from 8 species, was “replication, recombination and repair”. Four genes were annotated as transposases, 22 genes were from a family associated with plasmids (>10% of its members annotated as plasmids), and five genes were from a family associated with phages. With all of these annotations combined, in addition to the understanding these genes are significant beyond the overall population structure of their species, we conclude that many of the significant genes identified by the β test are likely to be components of mobile elements.

### Seven-Gene GRF operon is a structural variant in *Faecalibacterium prausnitzii*

One significant gene identified by microSLAM was annotated as “subunit D of the fructoselysine/glucoselysine phosphotransferase (PTS) system” by BlastKOALA [51] (S4 Table). It was negatively associated with IBD case status in *Faecalibacterium prausnitzii D* (UHGG species id 102272), a species whose relative abundance is positively associated with IBD in our compendium. This hit intrigued us, because *F. prausnitzii* is a well-studied bacteria with roles in short-chain fatty acid metabolism and inflammation [52,53]. Predicting a new molecular mechanism underlying this host-microbe interaction would enable future functional studies (e.g., in gnotobiotic mice) and potentially could be useful for developing diagnostics, dietary interventions, or other therapies.

To explore this gene family, we first compiled 85 high-quality and diverse *F. prausnitzii* genome assemblies from NCBI (S5 Table) and clustered them into eight clades (Fig 5A; Methods). We observed that seven genes (plus occasionally an eighth gene) were consistently found together, with a conserved order and orientation across 53% of the NCBI *F. prausnitzii* genomes (49/85) (S4 Fig). Annotations suggest that these genes encode a fructoselysine/glucoselysine PTS system operon. Having established that this operon is variably present across distantly related *F. prausnitzii* strains, we expanded our search to include all high-quality *F. prausnitzii* genomes available in the United Human Gut Microbiome Database (UHGG v2) [1]. This analysis indicated that the complete seven-gene operon is present in all nine *F. prausnitzii* clades, with between ∼3% and ∼24% of genomes per clade containing the operon (Table 1).

**Figure 5.**
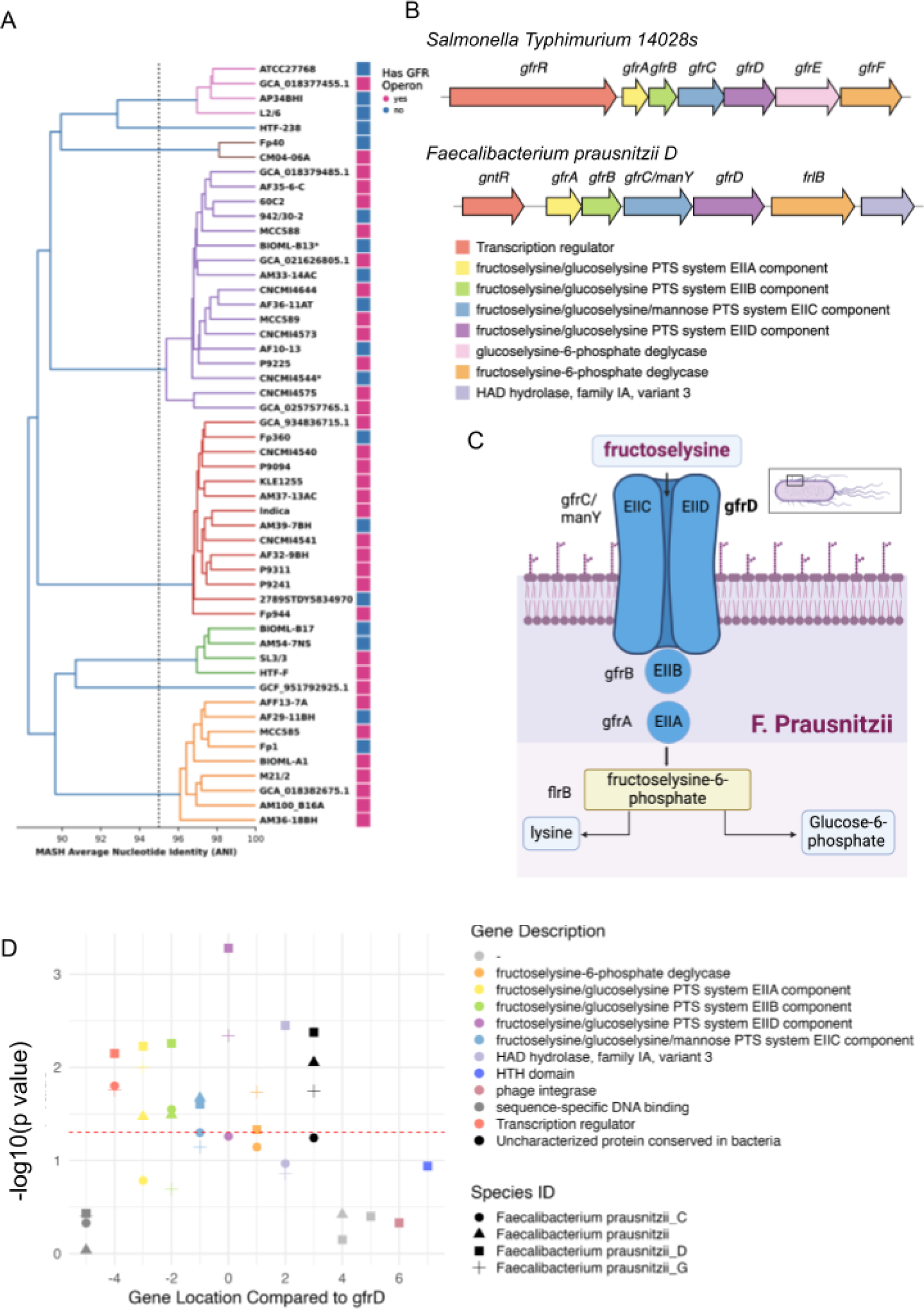
Investigation of *F. prausnitzii* fructoselysine PTS system operon. A) 52 representative genomes selected from NCBI and colored by the dRep secondary cluster (Selection of *Faecalibacterium prausnitzii* genomes, Methods). B) Comparison of *S. Typhimurium* operon to operon in *F. prausnitzii D*. C) Graphic of the *F. prausnitzii* fructoselysine PTS system operon and its products (made in BioRender). D) P-values for *F. prausnitzii* fructoselysine PTS system operon genes in microSLAM β tests across the four *F. prausnitzii* species defined by UHGG. The flanking genes are much less significant than the genes within the operon. Subunit D (most significant gene in microSLAM analysis) is located at 0, and all other indices are relative to this gene.

**Table 1.** A seven-gene operon present in all nine F. prausnitzii clades in UHGG v2.

| Species ID | Species name | Annotated as<br>plasmid | Not annotated<br>as plasmid | Total genome<br>counts | Proportion with<br>operon present |
| --- | --- | --- | --- | --- | --- |
| 100022 | *F. prausnitzii* C | 30 | 91 | 1258 | 0.096 |
| 100039 | *F. prausnitzii* H | 1 | 5 | 221 | 0.027 |
| 100195 | *F. prausnitzii* E | 6 | 4 | 244 | 0.041 |
| 101255 | *F. prausnitzii* F | 2 | 4 | 55 | 0.109 |
| 101300 | *F. prausnitzii* | 119 | 223 | 1446 | 0.237 |
| 102272 | *F. prausnitzii* D | 88 | 177 | 1280 | 0.207 |
| 102274 | *F. prausnitzii* I | 5 | 0 | 179 | 0.028 |
| 102545 | *F. prausnitzii* G | 124 | 127 | 2891 | 0.087 |
| 102619 | *F. prausnitzii* J | 25 | 32 | 236 | 0.242 |

Since genes can be syntenic without being functionally related, we conducted further analysis to determine the relationship between the seven-gene operon in *F. prausnitzii* and the well-characterized GFR operon in *Salmonella Typhimurium 14028s* [54]. We successfully mapped five genes from the *F. prausnitzii* operon to the corresponding *S. Typhimurium* operon (*gfrABCDF*) (Fig 5B). Notably, the *gfrE* gene, encoding a deglycase that cleaves glucoselysine 6-phosphate, is absent in *F. prausnitzii*. In addition, the operon in *F. prausnitzii* includes a gene without homology to any gene in the GFR *S. Typhimurium* operon. The regulatory genes, which are located at the start of the operons, also differ between the two species. We hypothesize the seven-gene *F. prausnitzii* operon identified in our analysis functions solely as a fructoselysine PTS system (Fig 5C).

Fructoselysine is a spontaneous product of Amadori rearrangements, and its presence in the human gut environment can promote the growth of bacteria capable of importing and using this carbohydrate as an energy source [55,56]. In *F. prausnitzii*, this is performed by a seven-gene operon encoding proteins that phosphorylate the substrate while transporting it across the bacterial cell membrane, making it available as a source of carbon. While only subunit D of this operon was significant in microSLAM’s gene test after accounting for multiple comparisons, all of the genes in the operon had unadjusted p-values less than 0.05 and flanking genes in the *F. prausnitzii* reference genome did not (Fig 5D). Variability in gene detection from shotgun metagenomics data is a likely source of the difference in significance for subunit D versus the other genes. MicroSLAM analysis for three other *F. prausnitzii* species in the IBD compendium did not yield any significant genes, primarily due to inadequate sample sizes, which restricts the statistical power of microSLAM’s β test. Nonetheless, the gene presence/absence matrices for these species are consistent with the operon being variably present and depleted in IBD cases.

Altogether, these results suggest that the genes in this *F. prausnitzii* PTS operon are co-evolving in terms of their presence/absence across hosts, potentially independently of neighboring genes, and that the presence of this operon is more common in healthy hosts. Our data also suggest that the fructoselysine PTS system operon could be a mobile genetic element. Supporting this possibility, many NCBI and UHGG contigs carrying this operon are predicted by geNomad [57] to be plasmids (Table 1). We also observe sequences associated with mobile elements and horizontal gene transfer (HGT) in the genomic context surrounding the operon. These are computational predictions only, and no plasmids have been previously reported in *F. prausnitzii*. We therefore checked for the operon in the other 70 species in our microSLAM analysis, detecting it in strains from two other phyla: *Gemmiger qucibialis* (species_id 103937) and *Faecalibacterium sp90053945* (species_id 103899). It is also known to be present in *Escherichia coli*, *Bacillus subtilis* and *Agrobacterium tumefaciens Ti plasmid* [56]. While not conclusive, these data are consistent with HGT. Regardless of the mechanism of acquisition or loss, the variable presence/absence of the operon across *F. prausnitzii* strains indicates that certain lineages of this species can acquire and utilize fructoselysine, thereby enhancing their adaptability and competitiveness in the dynamic gut ecosystem relative to strains without the operon.

## Discussion

In this paper we introduce microSLAM, a method that implements population structure-aware metagenome-wide association studies. Using a generalized linear mixed modeling approach, we are able to include information about the genetic relatedness of the microbiomes within diverse samples and to model the association of this population structure with host traits, while adjusting for other covariates. We focused on case/control study designs here, but microSLAM can also be applied to quantitative traits. In addition to testing if population structure itself (i.e., specific strains) are associated with a trait, microSLAM also includes a test aimed at identifying trait-associated genes that are evolving somewhat independently of strain lineages. Through realistic simulation studies, we demonstrated that microSLAM controls Type 1 error and has reasonable power in cohorts with more than one hundred samples. Compared to standard glm, microSLAM’s gene-level β test controls false positives much more effectively, especially in species with notable population structure. When there is a significant population structure as well as a subset of genes that are more related to the phenotype than the strain signal, we showed that microSLAM increases specificity compared to glm. By providing microSLAM as an open-source R package, we provide a new tool for researchers to probe microbiome-host interactions with strain- and gene-level resolution. We focused on applying microSLAM to the human gut microbiome to identify associations for IBD, but our R package is directly applicable to additional host traits and to additional environments where metagenomics data enables the computation of gene presence/absence matrices.

In this study, we put together a metagenomic compendium of IBD samples. Analyzing this data with microSLAM, we discover a wide variety of population structures within human gut metagenomes. We identified 49 species with a population structure related to IBD. In addition, after adjusting for population structure, 57 microbial genes are significantly enriched in healthy subjects, and 26 are enriched in IBD patients. From the genes enriched in healthy subjects, we identified a *F. prausnitzii* fructoselysine PTS system operon that is present in all clades of this species, but in only a minority of genomes within each clade, suggestive of being a mobile genetic element or other rapidly lost/gained structural variant. The presence/absence of this operon may confer distinct metabolic advantages to different strains, including the ability of carriers to utilize fructoselysine as an energy source [55]. The potential impact on human health could be significant, given that *F. prausnitzii* is one of the most prominent butyrate producers in the human gut [32,58,59]. This might also lead to a greater resilience of the gut microbiota, offering enhanced protection against pathogenic bacteria and reducing risk of chronic disease. Therefore, future work aimed at understanding the mechanisms through which *F. prausnitzi*i acquires and disseminates this seven-gene operon is not only key to comprehending microbial ecology but also crucial for potential dietary or probiotic therapeutic interventions targeting the microbiome.

There are several limitations to the microSLAM method and implementation. First, we estimate the GRM using the same gene presence/absence data that we then used to test genes for their trait-associations. This approach has been used with mixed modeling with human genetic data, and has been shown to reduce power compared to estimating the GRM with an independent set of markers (e.g., variants on other chromosomes) [60]. We explored a similar approach by using single nucleotide polymorphisms (SNPs) in core genes for GRM and random effect estimation in the β test. But we found that for almost all species in our IBD compendium, the SNP data generated a GRM that was very different from the gene-based GRM, and hence SNPs were not good markers for estimating the population structure in gene presence/absence. Perhaps this approach could work with more investigation into how to pick SNPs for GRM estimation or with a different GRM distance metric.

Second, as is the case with any meta-analysis, we used samples collected from many studies, located in a variety of geographic locations. Publicly available metagenomics data rarely includes detailed information about potentially confounding variables, such as diets and medical care. Hence, these important covariates are not accounted for in our models. This means that there is a chance some of the significant τ tests were related to an unmeasured variable that happened to be associated with strain genetic differences. Since each study included both cases and controls, and we did not observe any strong correlations between study and microSLAM’s random effect estimates, we do not consider this a major problem.

Nonetheless, with so little meta-data it is important to acknowledge that the strain-disease associations we detected across studies could be confounded by unmeasured variables (e.g., diet that selects for certain strains and alters IBD risk). In the future, when studies include more covariates, microSLAM can adjust for these just as we adjusted for age (the only consistently reported covariate) in this project. Beyond confounding, more complete metadata also would be helpful for understanding the capabilities of our method and for functionally interpreting our IBD findings.

Third, we do not find many individual significant genes within this study. This could be partially due to lack of power, especially for 55/71 species with ≤ 160 samples. A much larger dataset would increase our ability to find IBD associations for strains and genes. It would also enable separate modeling of associations for subtypes of IBD, which may have different microbiome signatures [61]. Our simulations suggested that most species did not have sufficient power for separate Crohn’s disease and ulcerative colitis models in our IBD compendium. We did investigate if any microSLAM discoveries were mostly driven by one subtype or the other, and we observed very few examples of this one significant gene cluster GUT_GENOME040547_00268 from species *Phascolarctobacterium faecium* was found to be positively associated with the disease but was only present in CD patients. In addition, there were three species with a significant τ where less than 1/3rd of the IBD patients are labeled as UC and there are less than 10 UC patients (*CAG-180 sp000432435, Ruminiclostridium E siraeum, and UBA11524 sp000437595*) meaning most of the signal is from CD alone in those species. While this finding could indicate that most associations are truly shared, it is more likely that we only had sufficient power to detect associations supported by both subtypes and that other subtype-specific associations remain to be discovered in the future with larger individual cohorts. As researchers move towards testing for strain and gene associations in studies with hundreds or thousands of samples, microSLAM’s improved specificity and controlled Type 1 error rate, as compared to glm, will be even more important.

Fourth, it is possible that some of our discoveries were driven by metagenomic reads from the wrong species or gene creating a false signal of gene presence (or absence). This cross-mapping is a frequent issue in read-mapping-based genomic analysis, especially for closely related species or highly conserved genes [62]. Hence, we recommend validating microSLAM’s gene test results with complementary data. For example, in our investigation of the PTS operon, we confirmed the operon structure and variable presence across strains using high-quality genome assemblies. This, plus the fact that this operon is predominantly found within *F. prausnitzii* and not widely distributed in other species, substantially alleviates concerns about cross-mapping in this analysis.

Finally, many of the genes we identified were not annotated, leading to difficulty completing in-depth analyses of significant genes across species. For example, we lacked power to perform functional enrichment analyses despite seeing several consistent trends, such as mobile elements being discovered in multiple species. More gene annotations would help with this problem, but annotation alone is not enough to confirm function. We view microSLAM as an important first step for proposing candidate causal strains and genes that should be performed upstream of in vitro and in vivo experiments to test hypothesized functions of the discovered strains and pathways. The ability of microSLAM to detect associations for species whose relative abundance is not correlated with host traits and to accurately disentangle associations of individual genes versus groups of strain-defining genes make it a useful new hypothesis generating tool for microbiome research.

## Methods

### Compendium of IBD/healthy case/control metagenomic studies

We compiled a total of 2625 publicly available paired-end shotgun metagenomic samples, sourced from five studies related to either inflammatory bowel disease (IBD) or the Human Microbiome Project (HMP2) and having an average read count greater than 20 million (accession numbers: PRJNA400072, PRJNA398089, PRJEB15371, PRJEB5224 and PRJEB1220). A stringent sample selection process was implemented to ensure (1) all samples included comprehensive metadata, such as disease status, age, and antibiotic usage; and (2) only one sample was selected per subject, considering that multiple time points could have been sequenced from the same subject. Specifically, for multiple time point samples from HMP2, we adopted the same selection criterion used by (Lloyd-Price et al., 2019) – selecting week 20 or greater for all subjects, the maximum read count for healthy subjects and the time point with the highest dysbiosis score for IBD patients. The first time point was chosen for the MetaHit project (Almeida et al. 2021; Nielsen et al. 2014; Li et al. 2014). Ultimately, a total of 710 samples met these criteria and were included for downstream analysis (S1 Table, S6 Table).

### Bioinformatics analysis

Preprocessing of the downloaded metagenomic sequencing libraries was performed using a QC pipeline that includes the following steps: (1) Adapter removal and quality trimming: adapter sequences were removed and low-quality reads were trimmed using Trimmomatic (v 0.39) [63]. (2) Human contamination removal: reads were aligned against the complete human reference genome (CHM13v2.0) [64] and a collection of 2250 genomes known to be contaminated by human sequences [65], using Bowtie2 (v 2.5.1) [66] to identify and remove human contamination. (3) Low-complexity read filtering: low-complexity reads were filtered out using BBduk (v 37.62) [67]. This step involved removing reads with an average entropy less than 0.5, with entropy k-mer length of 5 and a sliding window of 50 (parameters: entropy=0.5, entropywindow=0.5, entropyk=4). Additionally, reads shorter than 50 base pairs (bp) post-filtering were removed (parameters: minlen=50). (4) Quality reporting: a quality report of the cleaned-up reads was generated with FastQC (v 0.12.1) [68]. After preprocessing, samples with read counts lower than 1 million were removed, resulting in 710 high-quality samples retained for further analysis.

### Pangenome profiling using MIDAS v3

To determine which species within the 710 samples are both prevalent and sufficiently abundant for pangenome profiling, we implemented a two-step analysis using MIDAS v3 [11]. In the first step, we quickly scanned each sample to detect the presence of species by assessing the vertical coverage of 15 universal single-copy marker genes across 3956 distinguishable species in the UHGG v2 database [1]. In the second step, we adopted a whole-genome read alignment-based methodology to quantify the abundance of each species. This involved running MIDAS’s single-nucleotide variant (SNV) pipeline for species that meet specific criteria: a median marker coverage of at least 2X and at least 50% of the marker genes uniquely covered. These steps ensure that our whole-genome based species abundance estimation analysis is restricted to species with substantial coverage across their genomes (horizontal coverage > 0.4, vertical coverage > 5). We further excluded sample-species pairs where the ratio of genome-wide vertical coverage to single-copy marker gene coverage exceeded 4, which helps us to eliminate potential false positives caused by cross-mapping of reads among closely related species and conserved gene families. This stringent criterion also improves computational efficiency [62]. After implementing the aforementioned filtering, 71 species that were present in more than 60 samples and met the abundance criteria were selected for subsequent pangenome profiling analysis. There were 619 samples with at least one species present.

To perform pangenome profiling, we utilized the Genes module from MIDAS v3 [11], which features careful curation of the pangenome database and comprehensive functional annotation. Specifically, a single Bowtie2 index was built for all 71 species, and QC-ed paired-end reads for each sample were aligned to this index. Our analysis included only genes covered by at least 4 reads (--read_depth 4). Genes with an estimated copy number greater than 0.4 were classified as present (--min_copy 0.4). This threshold was selected based on exploratory data analysis and simulations previously performed by our lab. The resulting sample-by-gene presence/absence binary matrix was then used for subsequent association analysis with microSLAM, excluding core genes (absent in less than 10 samples) and rare genes (present in less than 30 samples).

### Statistical model for strain-trait and gene-trait associations while accounting for population structure in metagenomes

We present microSLAM, a 3-step modeling procedure for detecting within-species genetic variation associated with host biology (Fig 1A). The inputs to microSLAM, for a given species, are a *p* x *n* binary matrix of gene family presence/absence values for *n* host samples and *p* gene families, a 1 x *n* vector of trait values for each sample (binary or quantitative), and optionally a *q* x *n* matrix *X* of data for *q* covariates. The outputs are a measure of population structure (τ) with a permutation p-value and, for each gene family, a coefficient (β) measuring the gene’s association with the trait (e.g., log odds ratio for binary traits and logistic regression) with its local false discovery rate (localFDR) adjusted p-value [69]. Results from different species can be interpreted jointly to identify shared trends in trait-associations, such as enriched pathways (see below).

MicroSLAM fits generalized linear mixed effects models that account for the genetic relatedness of strains of a given species across hosts. In Step 1, an *n* x *n* sample genetic relatedness matrix (GRM) is computed from the gene presence/absence matrix. To do so, we create an *n* x *n* Hamming distance matrix and then transform this into relatedness using 1-distance. The GRM is used in Step 2 to test if the species’ population structure is associated with the trait, which would indicate that hosts with similar trait values tend to have similar strains.

For example, for a case/control study, this step aims to detect species where a subset of related strains confers risk. We call Step 2 the τ test, because population structure is modeled using a parameter τ. In Step 3, random effects estimated from the GRM are used to adjust for population structure in a model that is used to test gene families for associations with the trait beyond simply being present in trait-associated strains. We call Step 3 the β test, because a parameter denoted β is used to quantify gene-trait associations.

In Step 2 (τ test), microSLAM fits a generalized linear mixed model. The trait *y*_i_: *i* = 1,…, *1* is modeled as a function of any covariates *X* (with coefficients **α**) and random effects *b*_i_: *i* = 1,…, *1* that are estimated from the GRM. The link function f() is the identity function for normalized quantitative traits (linear regression) or the logit function for binary traits (logistic regression):

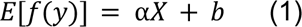

One key component of fitting Model 1 is estimating τ, the variance on the random effects, which depends on the association of the trait to the GRM. This is done iteratively using the average information restricted maximum likelihood (AI-REML) algorithm from the GMMAT [26] method. From this, we obtain a point estimate of τ, a point estimate of the random effects *b_i_*, and a statistic *T=b^2^/N*, that measures how associated the species’ population structure is with the trait. This T statistic is derived from [70] and computed using the linear setup from [71]. To assess the statistical significance of T, we randomly permute the trait values *B* times (e.g., *B*=1000), repeat model fitting, compute a T statistic for each permutation, and use these as an empirical null distribution to estimate a p-value based on how many of the permuted T statistics exceed the observed T statistic. Species with a significant T statistic have population structure that associates with the trait.

In Step 3 (β test), microSLAM fits a second model using the random effects (b) estimated in Step 2 and the presence/absence vector for each gene family, denoted *g* (with coefficients β):

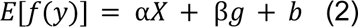

Model 2 is fit separately for each gene family within each species. β measures the gene’s association with the trait given the species’ population structure and the covariates. Similar to the strategy used in SAIGE [27], we directly calculate the score statistic for each gene by fitting the covariate and population structure adjusted genotype vector to the phenotype. Doing a direct computation given the random effect is an efficient strategy to reduce compute time; we only have to fit Model 1 once per species. Microbiome case/control studies are often unbalanced, for example, when a bacterial species is detected in many more controls than cases. To obtain accurate p-values in this scenario, we approximate the score statistics for testing the null hypothesis that β is zero using the Saddle Point Approximation (SPA) of the true distribution, as implemented in SAIGE.

To adjust the resulting p-values for multiple testing, we use localFDR [69], which accounts for the high correlation between gene families (i.e., when genes co-occur across strains) that invalidates methods such as Benjamini-Hochberg FDR [72] or Storey’s q-value [73]. We transform SPA p-values into Z-values by dividing by two, multiplying times the sign of the estimated β coefficient, and converting the resulting numbers to quantiles. Then, localFDR uses maximum likelihood estimation to approximate the null Z-value distribution and identify Z statistics that deviate from this distribution. We implement this using the *locfdr* v1.1-8 package in R, fitting the null distribution to the Z-values between the 10th and 90th percentiles across all species (Fig S5).

### Generalized Linear Model

A standard generalized linear model (glm) [74] was fit for all genes for all species that were analyzed with microSLAM. This is a logistic regression model using *glm* in R with case/control status as the outcome, gene presence/absence as a predictor, and age as a covariate.

### τ Test Simulations

We performed simulations to assess the false positive rate and power of microSLAM’s τ test. To assess the false positive rate (simulation 1), we set up a simulation where a binary trait was generated independently of GRMs. For each of 1000 iterations and n=100 samples, the trait *y* was simulated using a binomial distribution with a success probability of 0.5, and a covariate was simulated with a normal distribution centered at 45 with a standard deviation of 15, similar to the age distribution in our IBD compendium. Next, for each iteration, a gene presence/absence matrix was simulated with p=1000 genes. This included 400 “core” genes simulated from a binomial distribution with a success probability of 0.8, 400 “accessory” genes simulated from a binomial distribution with a success probability of 0.2, and 200 genes simulated based on presence of a strain unrealted to the trait *y*, as follows. The strain’s presence/absene across samples was simulated using a binomial distribution with a success probability of 0.5, and then presence/absence for each of the 200 genes was set to absent if the strain was absent and simulated from a binomial distribution if the strain was present, where the success probabilities were chosen such that the average odds ratio of a given gene being present if the strain is present is 1.8. After the genes are simulated, the GRM is calculated and the population structure test is run with 100 permutations. The p-value is calculated for each iteration as the number of permutations with a more extreme T statistic than the observed T statistic. The false positive rate is calculated as the number of iterations with a p-value <0.05.

The power test (simulation 2) is carried out in a similar fashion, with two key changes. First, we explored a range of sample sizes (n=60, 100, 250) to assess the relationship between sample size and power. Second, we simulated the trait y based on presence/absence of the simulated strain. Specifically, we explored a range of effect sizes, quantified with an odds ratio θ/(1 − θ) ranging from 1.0 to 2.5. For a given odds ratio, we set *strain_δ_ = (1 − strain) * (1 − θ) + strain * θ* and then generated the trait *y*_strain_ using a binomial distribution with success probability equal to strain_δ_: *y*_strain_ = rbinom(*n*, 1, strain_δ_). This creates a stronger relationship between the trait and presence of the strain as the odds ratio increases. For each odds ratio and sample size, 125 iterations were run and power was calculated as the proportion of iterations with a significant τ test divided by 125.

### τ Test Simulations

We performed simulations to assess the false positive rate and power of microSLAM’s β test versus a standard glm. To assess false positives (simulation 1), we generated data in which no genes were associated with the trait, so that all genes are false positives. We computed a p-value for each gene using each modeling approach and tracked the proportion of genes with p<0.05. In order to simulate real population structure, while introducing some random variation, we used the observed GRM for each of the 71 species in our metagenomic compendium to help generate simulated gene presence/absence matrices. Specifically, we first decomposed the observed GRM for each species into its first 10 principal components (PCs). We then standardized each PC by dividing each value by the PC’s standard deviation pc_std_ = pc/sd(pc) and computed the standard normal probability for each sample’s loading on each standardized PC (one per sample *i* per PC dimension *j*): *p_i,j_* = pnorm*_std_*). The probabilities *p_i,j_* retain relationships between samples across the 10 dimensions. For each of the 10 PCs, we simulated the presence/absence of 90 genes using a binomial distribution with a success probability equal to the sample’s *p_i,j_* for that PC, for a total of 900 genes correlated with one dimension of the population structure. We also simulated 100 uncorrelated genes using a binomial distribution with a success probability chosen from a uniform distribution between 0.2 and 0.8. From the resulting 1000 x n gene presence/absence matrix, we simulated a binary trait (*y)* using the first two PCs (PC1 and PC2), as follows. We set *y* equal to one in a given sample if its loadings on PC1 and PC2 had opposite signs (either PC1>0 and PC2<0 or PC1<0 and PC2>0). This created a nonlinear relationship between *y* and 180 of the simulated genes (Fig S6).

For each simulated *y* and gene presence/absence matrix, we ran microSLAM to compute a GRM and estimate population structure (τ). These new τ’s were different from the species’ observed τ values and greatly varied across species, as in the observed data (Fig S7). After the GRM was calculated, and the τ test was run, both glm and microSLAM’s β test were run to test for gene-trait associations. The false positive rate was determined by summing the number of genes with p-values < 0.05 divided by the total number of genes, excluding genes simulated from PC1 or PC2 (p=820 genes).

To assess power for the β test (simulation 2), we start with the data from simulation 1 and set *y*_δ_ = (1 − *y*) * (1 − θ) + *y* * θ for an odds ratio of θ/(1 − θ). Then, we generated presence/absence for 100 additional genes from a binomial distribution with success probability *y*_δ_: *G*_*y* = rbinorm(n, *1*, *y*_δ_). These genes G_y are positives (associated with y) and all other genes are negatives (independent of y). At θ = 0. 5 (i.e., an odds ratio of one), the generated genes *G_y* will not be associated with the trait. At θ = 0. 55, the average odds ratio will be 1.2. We investigated θ values between 0.52 and 0.78. We checked, and the odds ratios across simulations with the same θ value did not deviate more than 0.1 from the expected values.

We ran glm and microSLAM on each simulated dataset. As expected, the population structure test yielded estimated τ values that increase notably with the simulated odds ratio (i.e., as the association between y and the genes *G_y* increases). In order to assess power, we used the negative genes to establish a significance threshold for each modeling approach such that the empirical false positive rate would be no more than 0.05. Applying these thresholds to the positive genes *G_y*, we computed power as the proportion of positive genes detected. Power was compared between glm and microSLAM across odds ratios and species (each with different sample sizes and GRMs).

### Relative abundance test

We calculated the relative abundance of each species by downloading the UHGG v2 kraken database from MGnify [75] and running Kraken2 [45,47] with options *–paired--minimum-hit-groups 3* and then bracken [45,46] with options *-l S-t 1000.* We computed relative abundance as a given species’ bracken coverage divided by the total coverage, and we removed species with less than 0.05% relative abundance. We then performed logistic regression, using the case/control label as the dependent variable (y) and relative abundance as the independent variable, with age as a covariate. The estimated log odds ratios and p-values from these logistic regression analyses were compared with outputs from the microSLAM τ test, after using localFDR to adjust p-values at a 0.1 level.

### Identification of seven-gene GFR operon in *F. prausnitzii*

The gene *grfD* from *F. prausnitzii* (UHGG species id 102272) reported as significant by microSLAM’s β test corresponds to the EIID component and is part of a putative GFR operon that encodes the Fructoselysine PTS system. A similar gene in *S. Typhimurium 14028s* has been identified as responsible for the utilization of fructoselysine [54,76]. To determine whether this EIID gene is part of a gene cluster that forms an operon - that is, genes that are sequentially arranged on the chromosome and co-regulated - we conducted the following analysis. First, we retrieved the neighboring genes upstream and downstream of this EIID gene in the UHGG reference genome for this *F. prausnitzii* species, considering up to five genes in each direction. We then used blastn [77] to identify homologous regions in three different sets of genomes: (1) 85 NCBI *F. prausnitzii* genomes, (2) all genes in the nine UHGG *F. prausnitzii* species clusters including metagenome-assembled-genomes (MAGs), and (3) all MAGs in the 71 species that were investigated in this study. These five genes from *F. prausnitzii* were also aligned to the corresponding *S. Typhimurium* operon (gfrABCDF) using Blastp [77]. All genomes were annotated using Prokka [78]. Genes were annotated with BlastKOALA [51] and eggNOG-mapper [79].

### Selection of *Faecalibacterium prausnitzii* genomes

To avoid incorrectly assessing the seven-gene GFR operon as incomplete in an assembly simply due to fragmented contigs, we only selected high-quality *F. prausnitzii* genomes with assembly levels of scaffold, chromosome, or complete genome, and we specifically excluded atypical genomes. In total, we downloaded 105 genomes of *F. prausnitzii* from NCBI (using the taxon identifier 853). We assessed the genome quality using CheckM [80], and retained only genomes that met the following criteria: completeness >= 90, contamination <=5, and strain heterogeneity <= 10. After this filtering, 85 *F. prausnitzii* NCBI genomes were retained for the GFR operon screening analysis (Fig S4). Next, we used dRep [81] to perform pairwise genome comparisons based on Average Nucleotide Identity (ANI). This dRep analysis involved first clustering all the genomes using the Mash heuristic for ANI [82] and subsequently using MUMMER [83] to compute ANI on sets of genomes that have at least 90% Mash ANI before performing a secondary clustering. As a result, 52 secondary clusters were formed at 98% MUMMER ANI (-comp 90-con 5-pa 0.95-sa 0.98-nc 0.8). Hierarchical clustering of the 52 representative genomes using average linkage was performed using the pairwise MASH similarity matrix (‘scipy.cluster.hierarchy’ package).

In addition to NCBI Genomes, we also collected all nine *F. prausnitzii* species clusters from UHGG v2 [1], using the same selection criteria to ensure assembly quality. We calculated the pairwise genome similarity between the resulting 52 *F. prausnitzii* genomes and the nine representative genomes from UHGG *F. prausnitzii* species using fastANI [84]. We also compared each NCBI *F. prausnitzii* genome to the nine representative genomes from UHGG, and each NCBI genome was assigned to the UHGG species cluster with the highest ANI (ANI >= 95%). If this similarity level was not reached, the NCBI genome remained unassigned. Eight *F. prausnitzii* species in UHGG were represented by the 52 NCBI *F. prausnitzii* genomes. The information for the 85 genomes is available in Supplemental Table 5.

## Acknowledgments

We would like to thank Byron Smith, Abigail Lind, Xiaofan Jin, Lei Liu, and Eran Segal for their insightful discussions about the project.

## Code reporting

The microSLAM R package is available at https://github.com/miriam-goldman/microSLAM.

## Data availability statement

Reference genomes and metagenomic data analyzed for this study are available in public repositories as described in the Methods.

## Supplemental Materials

### Supplemental Tables

**Supplemental Table 1 - IBD compendium data information**

**Supplemental Table 2 - IBD compendium species results**

**Supplemental Table 3 - IBD compendium gene family results**

**Supplemental Table 4 - PTS operon Blast KOALA results**

**Supplemental Table 5 - NCBI *F. prausnitzii* genomes**

**Supplemental Table 6 - Quality control for all samples**

## Supplemental Text

### Genetic Relatedness Matrix

We used the pairwise Hamming distance based on the gene presence/absence matrix to create the Genetic Relatedness Matrix (GRM) for a set of samples. For *N* genes where A_i_ and B_i_ are a binary presence or absence of gene *i* in sample A and in sample B, respectively, we define genetic similarity as 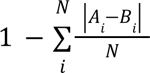. This is computed for all pairs of samples where a given species is present to create that species’ GRM ψ.

If wanted, one could instead use a single-nucleotide variant (SNV) presence/absence or frequency matrix instead of a gene presence/absence matrix to compute the GRM. For SNV distance, we recommend using the Manhattan distance. The resulting GRM based on the set of bi-allelic polymorphisms genotyped by MIDAS v3 or another tool would approximate the average nucleotide identity (ANI) between samples. We explored this approach and found that polymorphism based GRMs were generally very different from gene presence/absence based GRMs for the same species. In simulations, this led to higher false positive rates for the microSLAM β test (similar to glm), presumably because there was trait-associated population structure in the gene presence/absence matrix for which this approach did not fully adjust. Further investigation into the selection of SNVs or distance metric could potentially make this strategy more effective.

### Microbiome generalized linear mixed model for binary traits: additional details about microSLAM’s modeling approach

In a case-control study with sample size N, we denote the status of the *i*th individual with **y*_i_*= 1 or 0, depending on whether it is a case or a control. Let the 1 × (1 + *p*) vector *X_i_* represent *p* covariates, plus an intercept term, and let *G_i_* represent the presence or absence of a gene; this can also be replaced with the copy number of a gene, as estimated by MIDAS v3 or other tools. The logistic mixed model can be written as:

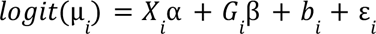

where µ*_i_* = *p*(*y_i_*= 1|*b_i_*, *G_i_*, *b_i_*) is the probability that the *i*’th individual is a case given the covariates, gene presence/absence vector, and the random effect *b_i_* that is estimated by microSLAM. The random effect *b_i_* is modeled τ(0, τ ψ) where ψ the N×N GRM described above, and τ is the estimated additive genetic variance. The *var*(*y_i_*| *b_i_*) = ϕ var(µ*_i_*), in the case of a binary trait the random parameter Φ=1. The parameter α is a 1 × (1 + *p*) coefficient vector of fixed effects and β is a coefficient representing the log odds ratio for the association between the gene’s presence and the trait. For a quantitative trait, *y_i_* is a real number and the model is a linear mixed model rather than logistic, so that β represents the expected change in the trait for the gene being present versus absent. Everything else is the same.

### Estimating the coefficients and variance component: fitting microSLAM for each species

We employ the same restricted log-likelihood and average information matrix for estimating the coefficients and variance components as were employed in GMMAT [26] and Saige [27]. For more details on deriving these estimation procedures, refer to those studies and to Clayton and Breslow [25]. We also follow Saige’s multi-step process to estimate the random effects and then use these in the logistic (or linear) model presented in the previous section. This helps us in two ways: 1) it reduces computational time significantly as random effects only have to be estimated one time for each species (not once for every gene in every species), and 2) avoiding refitting the random effect for every gene provides a more robust estimate.

Unlike Saige, we do not use PCG, randomized trace estimator, or a low-rank GRM. These are designed to reduce computation and memory costs within the context of human genomes with millions of genetic variants, but these are not major problems for us given the size of the datasets in this study. Also, our GRMs are naturally full-rank. These computational shortcuts could be implemented if needed.

### Score testing for the GRM: τ test modeling

We detail microSLAM’s τ test, a new statistical procedure to inform the user whether the species’ GRM is significantly related to the trait. This would indicate that a subset of related strains can predict the trait. We consider random effects *b_i_* ∼ τ(0, τψ), as described above, then compare the models:

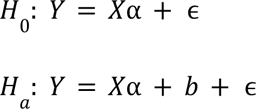

After the models have been fit (estimation converges), we have 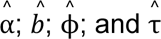. We also compute a working vector

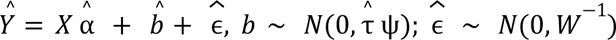

The test statistic for the τ test can be written as:

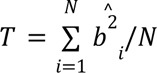

This is the sample variance of the estimated random effects *b_i_*. This statistic involves the sum of the squared random effect estimates. The null hypothesis is that T=0 (i.e., the random effects do not help to explain variation in the trait). To compute a p-value for T without making assumptions about its distribution, we use a permutation test.

### Score testing for gene presence/absence: β test modeling

After we have fit the model described above for the τ test, we have estimates of the fixed effect coefficients 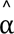, the random effects 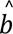, and the variance component parameters, 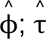. Using these, we construct a score test for each gene with the null hypothesis *H*_0_: β = 0. Suppose *G* is a τ × 1 genotype vector (where τ is the number of samples). 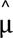 are the probabilities of the samples having the trait (e.g., being cases) given the covariates *X* and the random effects 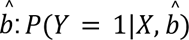. Let 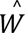 be a diagonal vector with elements 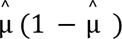 and 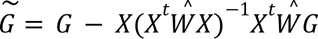 is the covariate-adjusted genotype vector. With 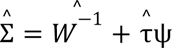 and 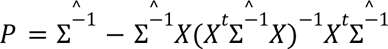 and a working vector 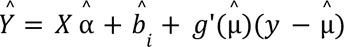, the score test statistics, assuming 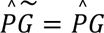 is:

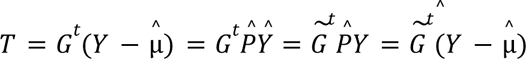

The variance of T is:

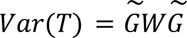

We estimate this directly for each gene *G*. As shown in [27] this is approximately equivalent to 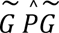 but much faster to compute, plus the approximation is conservative.

The effect size 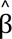 is the natural log of the odds ratio. We can estimate this using the variance component estimate under the null hypothesis.

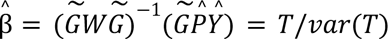

The standard error of 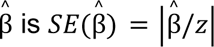 where *z* is the z-score corresponding to the p-value divided by 2.

### Simulations

For the β test simulation 3, we sought to generate gene presence/absence matrices, trait values, and age values without using any real sample data. First, we simulated a trait from the binomial distribution with a success probability of 0.5. We assume that there are two strains, one that is correlated with the trait (*odd ratio = 2. 22*) and one uncorrelated with it. For the correlated strain, we simulated 300 genes at a presence level of 0.5 and an odds ratio of 4.0 for the gene being present given a person has the strain. We simulated one-third of the samples having the uncorrelated strain and modeled this with 250 genes with low binomial presence rates (*p* = 0. 3) and an odds ratio of 4.0 for the gene being present given a person has the strain. Then we modeled 300 genes that were “core” across all samples; these were drawn from a binomial with a success probability of 0.8. We additionally simulated 150 genes that were non-strain associated “accessory” genes; these were drawn from a binomial success probability of 0.2. Last, we simulated at least one gene (*G_y_*) that is even more highly correlated with the trait than is the correlated strain (*odd ratio* = 2. 44). The more genes in *G_y_* (we investigated 1, 2, or 3 genes) and the stronger the relationship between *G_y_* and the trait, the higher the parameter τ will be. The resulting gene presence/absence matrices naturally have a range of different values of τ. Age was randomly generated with parameters similar to the IBD data: *ceiling(rnorm(N, mean = 45, sd = 15))*. We repeated Simulation 3 with the number of samples varying from 60 to 250.

---

**Supplemental Figure S1.**
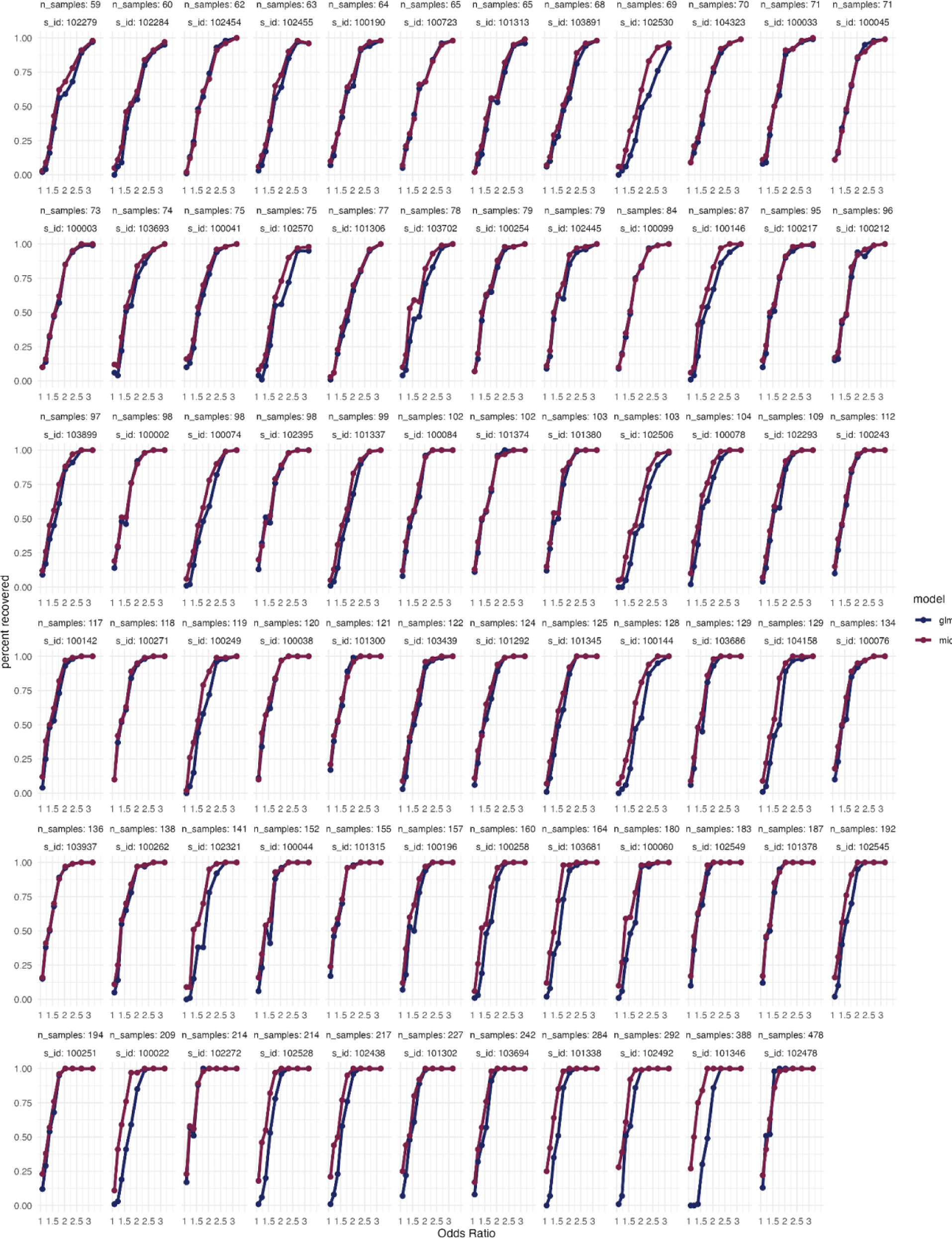
GLM and microSLAM β test power evaluations for 71 simulated species. These plots show estimated power of β tests using data from simulation 2 in which a gene presence/absence matrix and binary trait were simulated based on the observed GRMs from the 71 species in the IBD compendium using a range of different effect sizes (odds ratios, horizontal axes). There is one panel per GRM (labeled with species ID), and panels are ordered from lowest to highest sample size.Power was computed as the proportion of positive genes discovered at an empirical localFDR of 0.05 for both microSLAM (*red*) and glm (*blue*). As the number of samples increases there tends to be a larger difference between the glm and the microSLAM models.

**Supplemental Figure S2.**
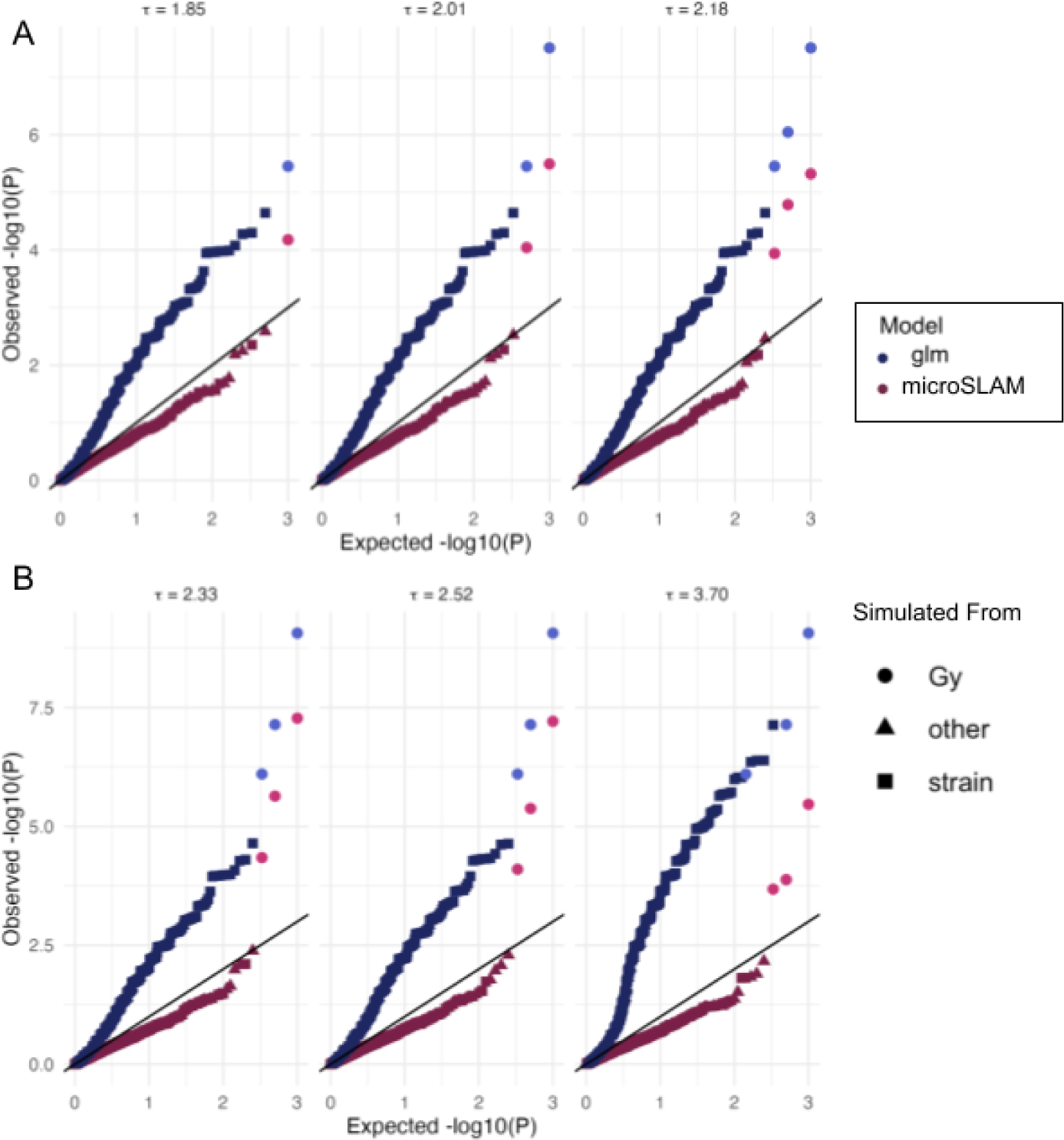
P-values from microSLAM’s β test are somewhat conservative, while GLM’s are inflated. Q-Qplots of β Test results for a simulation with positive genes (*G_y;_ pink circles*: microSLAM, *light blue circles*: glm) plus negative genes that are linked to a strain (strain; squares) or randomly generated (other; triangles) (β test simulation 3, **Supplemental Text).** A) Compared to glm (blue), microSLAM (red) better distinguishes the positive genes *G_y_* from those simulated from the strain. The number of positive genes was one (*left*), two (*middle*), or three (*right*). The value of τ increases with each additional gene *G_y_*. B) As the relationship between the strain and *y* is increased (left to right), the value of τ increases, and the rate of inflation increases for glm. Across different values of τ, microSLAM remains slightly conservative and continues to rank the positive genes *G_y_* highest, indicating high specificity.

**Supplemental Figure S3.**
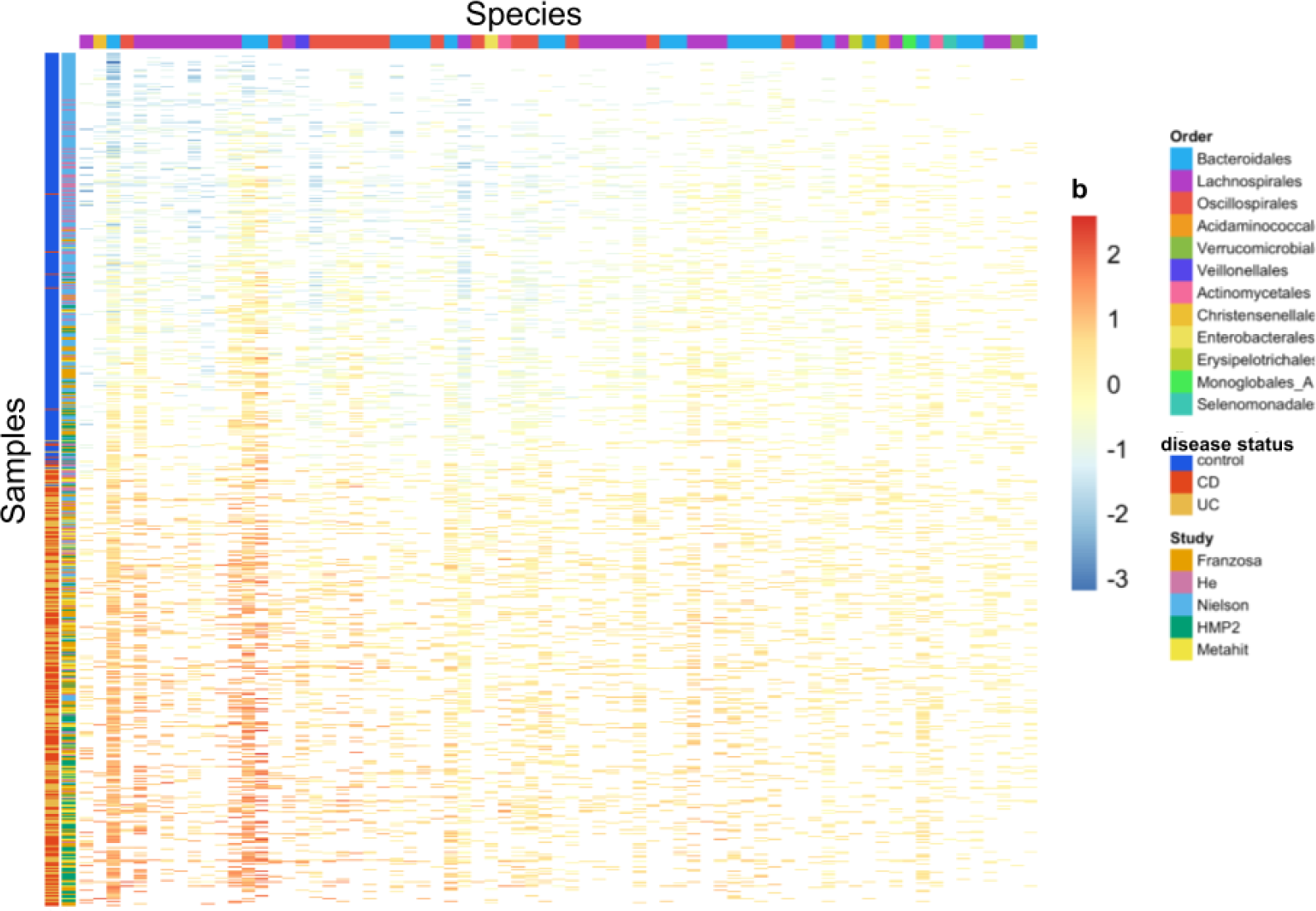
Random effect (b) values across studies and species. This heatmap shows microSLAM estimates of the random effect parameters (b) for each of 71 species across all samples where it was detected in the IBD compendium. The red-to-blue color scale denotes the association between strains and IBD (binary case/control status). Red: strains positively associated with IBD; Blue: strains negatively associated with IBD. The study and IBD subtype of each sample are shown on the left. IBD subtypes: Blue=control, red=Crohn’s disease (CD), yellow=Ulcerative colitis (UC). CD and UC were combined as cases in the microSLAM modeling. Studies: Franzosa (NCBI BioProject PRJNA400072; orange), He (PRJNA398089; pink), Nielsen (PRJEB15371; blue), HMP2 (PRJEB5224; green), MetaHIT (PRJEB1220; yellow). Species are ordered by the standard deviation of *b* (left=highest standard deviation), where higher standard deviation indicates greater strain diversity that is associated with case/control status. The samples in each column are ordered by lowest to highest average b value. A few studies (e.g., Nielsen) have more controls than others, but there is no systematic relationship between study and population structure. CD and UC tend to have similar distributions of *b* values (i.e., red and yellow are mixed on the left side bar). While we cannot rule out confounders that were unmeasured in the publicly available data that we could access, these patterns suggest that our findings are not obviously biased by differences in study population (e.g., diet, medical care, geography, type of IBD) that could confound measured associations between case/control status and microbiome strains and genes.

**Supplemental Figure S4.**
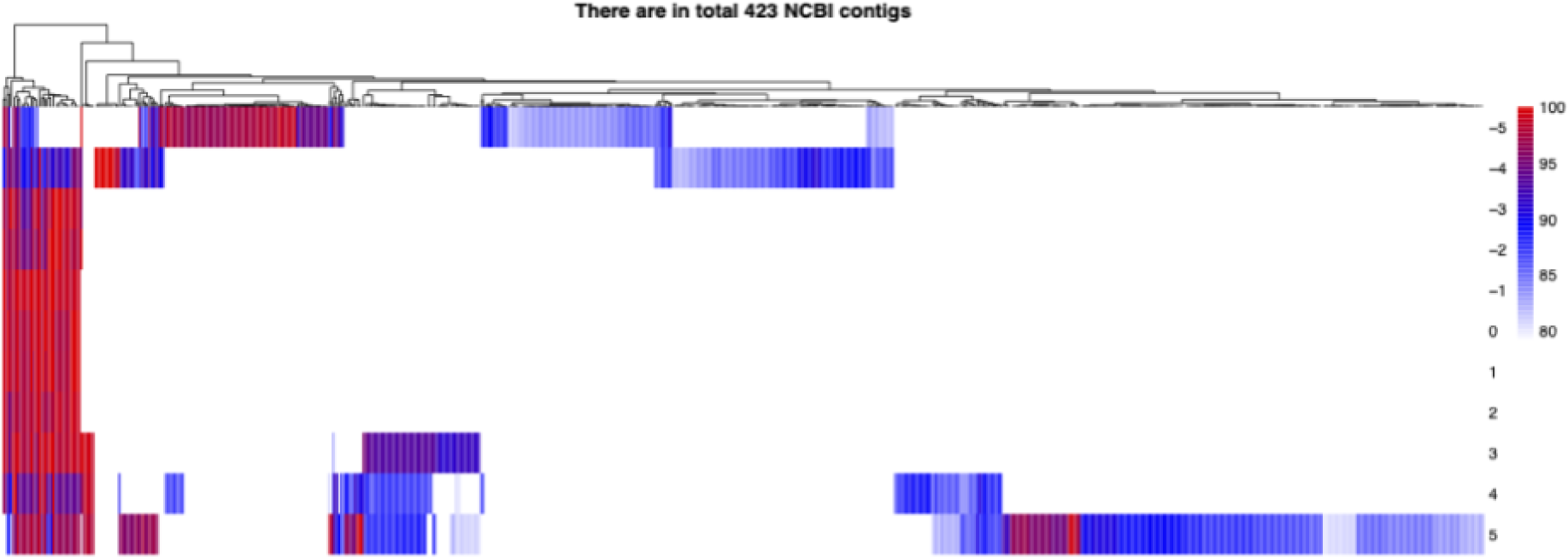
*F. prausnitzii PTS* operon evolves as a unit across diverse genomes. This operon comprises seven genes (occasionally eight genes) that were consistently present or absent together across 53% (49/85) of *F. prausnitzii* genomes from NCBI. The order and orientation of genes in the operon is conserved. This heatmap shows the genes (rows; position 0 is *gfrD,* which was significant after localFDR adjustment of microSLAM β test p-values). The other genes were significant before localFDR adjustment and are indexed relative to *gfrD* in the heatmap. Columns represent 423 contigs from 85 *F. prausnitzii* high-quality NCBI genomes. The color of the heatmap shows the blastn sequence similarity of the gene sequence in the contig compared to the sequence in the *F. prausnitzii* reference genome used in our microSLAM analysis (red=highest similarity, white=no significant match). The seven genes in the operon (middle rows of the heatmap) have high sequence similarity when they are present and are present together (red on left), whereas flanking genes are more variably present and have lower sequence identity (blue in top and bottom rows).

**Supplemental Figure S5.**
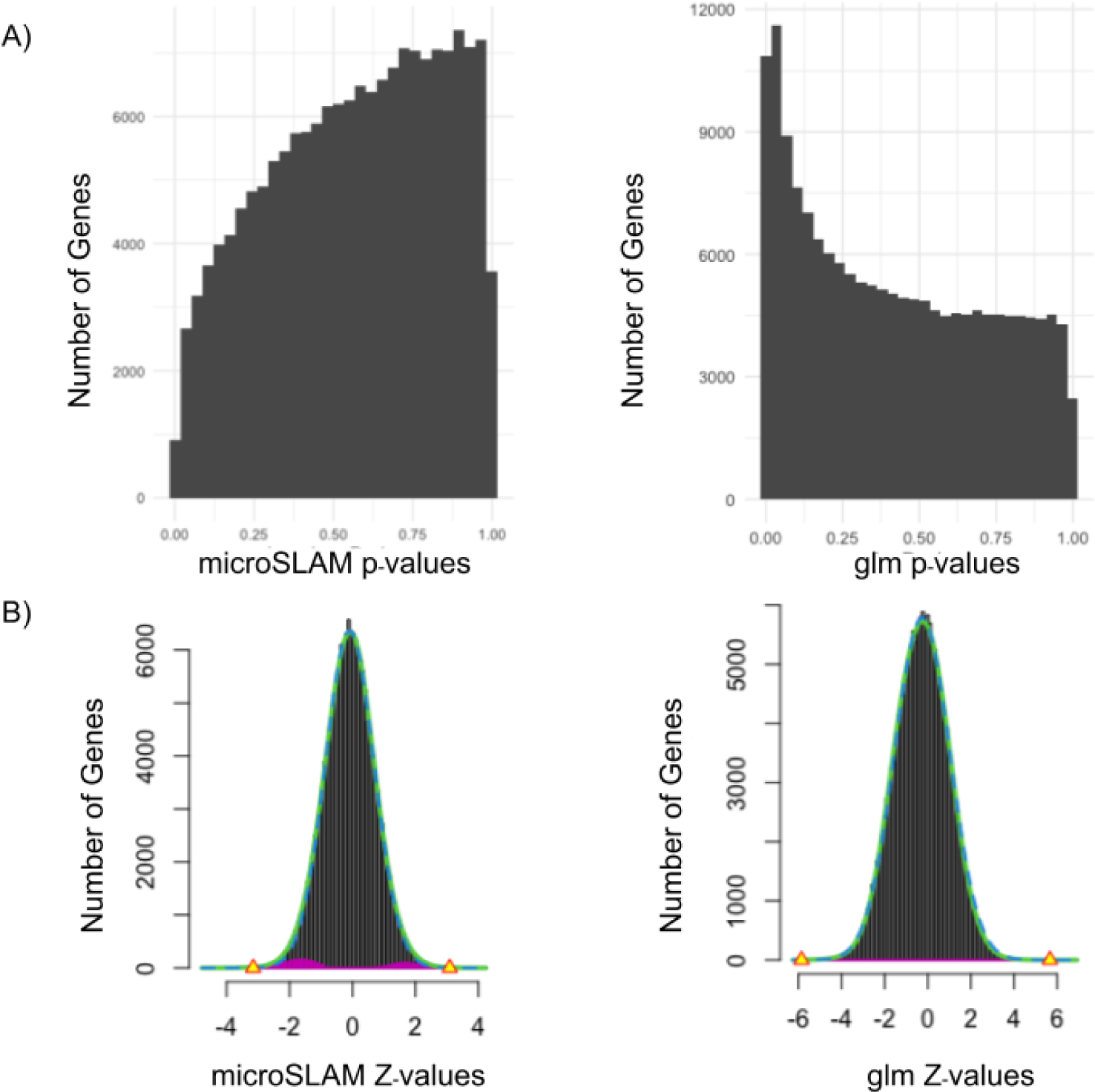
LocalFDR p-values and Z-values. A) Histogram of p-values for microSLAM’s β test (left) and glm (right). B) Output from localFDR showing the distribution of the null z-values (green) versus the distribution of the z-values that do not follow the null (pink). Yellow triangles denote the z-value thresholds corresponding to a localFDR of 0.2.

**Supplemental Figure S6.**
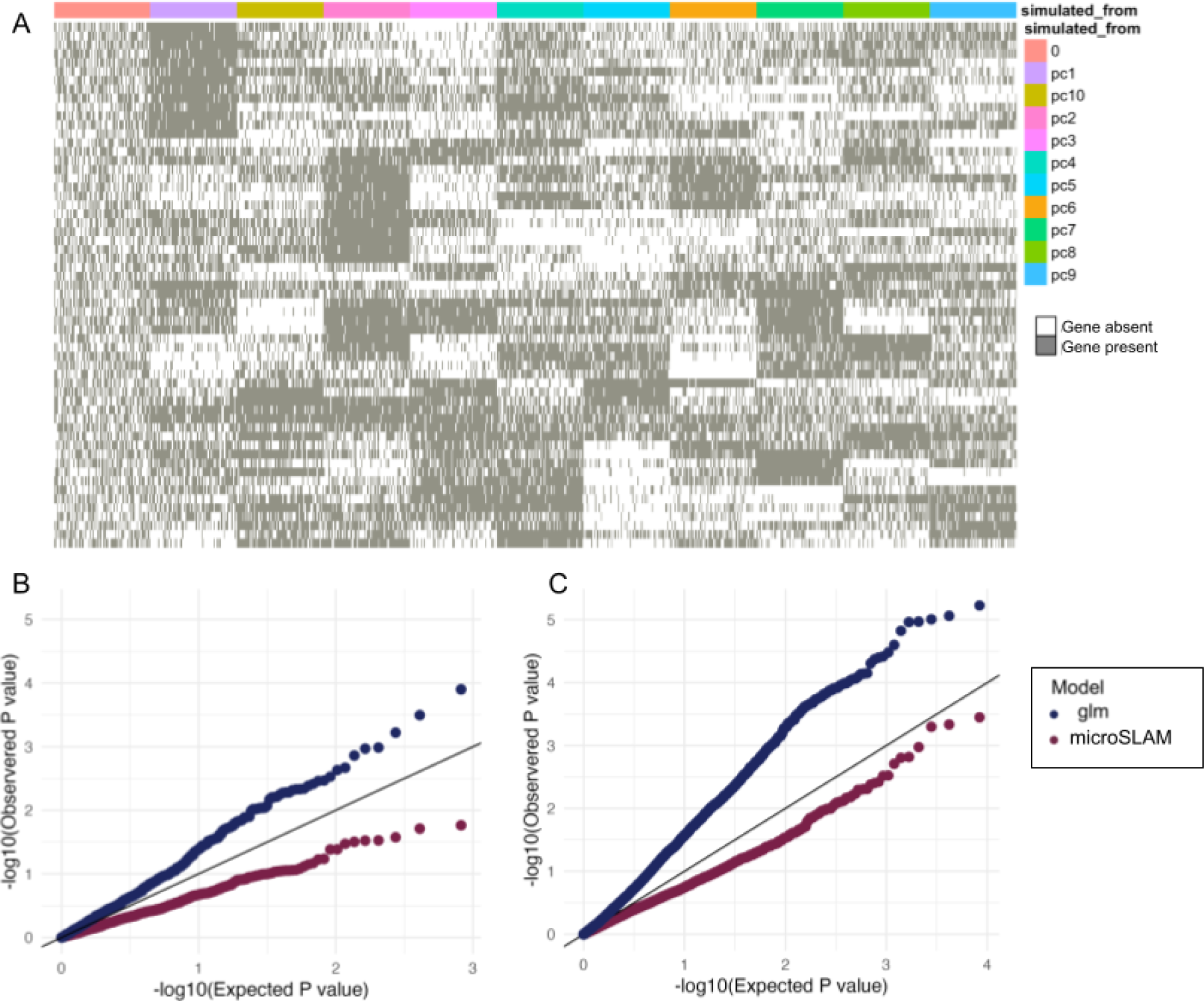
Example of data and results from microSLAM τ test Simulation 1. A) Simulated gene presence/absence matrix based on the GRM of *Bacteroides thetaiotaomicron* plotted as a heatmap (grey=gene present in a given sample, white=gene absent). Genes are in columns and are labeled according to how they were simulated (0=random, pc1-10=using one of the first 10 principal components of the observed GRM for *B. thetaiotaomicron*. This presence/absence matrix has a some population structure (estimated τ = 2.30), but no genes were simulated to be associated directly associated with the trait which is defined by the first two PCs. B) Q-Qplot of p-values from all genes not from PC1 or 2 from microSLAM’s τ test (red) and glm (blue) applied to the simulated gene presence/absence matrix in (A). There is a much higher error rate for the glm model. On the other hand, microSLAM is overly conservative (i.e., underpowered). C) Q-Q plot for microSLAM’s τ test (red) and glm (blue) applied to the observed *B. thetaiotaomicron* gene presence/absence matrix from the IBD compendium. The trends are very similar to those in the simulation.

**Supplemental Figure S7.**
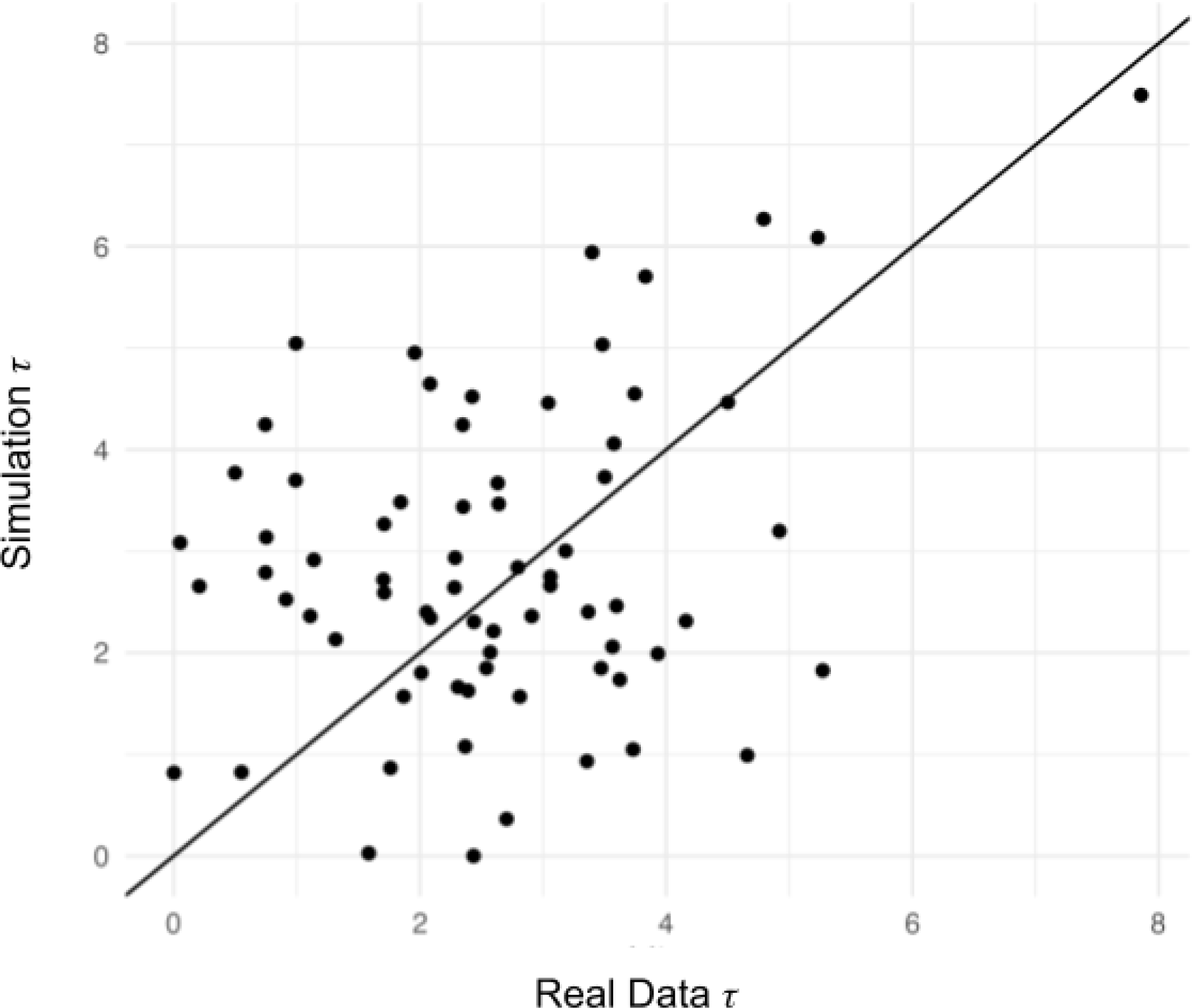
τ Test simulation τ values versus observed τ values in IBD compendium. In the β Test Simulation 1 and 2 set up, we generated gene presence/absence matrices using the observed GRMs for the 71 species in the IBD compendium. Our objective was to generate simulated data that was similar to but not identical to the observed data (**Methods**). This scatter plot shows the τ values estimated by microSLAM on the simulated data (*y axis)* compared to the corresponding τ values estimated from the real data in the IBD compendium (*x axis*). The n τ from the simulation cover a similar range of values as those from the real data while not being highly correlated.

## References

1. Almeida A, Nayfach S, Boland M, Strozzi F, Beracochea M, Shi ZJ, et al. A unified catalog of 204,938 reference genomes from the human gut microbiome. Nat Biotechnol. 2021;39: 105–114. doi:10.1038/s41587-020-0603-3

2. Kurilshikov A, Medina-Gomez C, Bacigalupe R, Radjabzadeh D, Wang J, Demirkan A, et al. Large-scale association analyses identify host factors influencing human gut microbiome composition. Nat Genet. 2021;53: 156–165. doi:10.1038/s41588-020-00763-1

3. Wang J, Thingholm LB, Skiecevičienė J, Rausch P, Kummen M, Hov JR, et al. Genome-wide association analysis identifies variation in vitamin D receptor and other host factors influencing the gut microbiota. Nat Genet. 2016;48: 1396–1406. doi:10.1038/ng.3695

4. Lopera-Maya EA, Kurilshikov A, van der Graaf A, Hu S, Andreu-Sánchez S, Chen L, et al. Effect of host genetics on the gut microbiome in 7,738 participants of the Dutch Microbiome Project. Nat Genet. 2022;54: 143–151. doi:10.1038/s41588-021-00992-y

5. Falony G, Joossens M, Vieira-Silva S, Wang J, Darzi Y, Faust K, et al. Population-level analysis of gut microbiome variation. Science. 2016;352: 560–564. doi:10.1126/science.aad3503

6. McCarthy AJ, Loeffler A, Witney AA, Gould KA, Lloyd DH, Lindsay JA. Extensive horizontal gene transfer during Staphylococcus aureus co-colonization in vivo. Genome Biol Evol. 2014;6: 2697–2708. doi:10.1093/gbe/evu214

7. McDougal LK, Steward CD, Killgore GE, Chaitram JM, McAllister SK, Tenover FC. Pulsed-field gel electrophoresis typing of oxacillin-resistant Staphylococcus aureus isolates from the United States: establishing a national database. J Clin Microbiol. 2003;41: 5113–5120. doi:10.1128/JCM.41.11.5113-5120.2003

8. Alkan C, Coe BP, Eichler EE. Genome structural variation discovery and genotyping. Nat Rev Genet. 2011;12: 363–376. doi:10.1038/nrg2958

9. Wang D, Doestzada M, Chen L, Andreu-Sánchez S, van den Munckhof ICL, Augustijn HE, et al. Characterization of gut microbial structural variations as determinants of human bile acid metabolism. Cell Host Microbe. 2021;29: 1802–1814.e5. doi:10.1016/j.chom.2021.11.003

10. Zeevi D, Korem T, Godneva A, Bar N, Kurilshikov A, Lotan-Pompan M, et al. Structural variation in the gut microbiome associates with host health. Nature. 2019;568: 43–48. doi:10.1038/s41586-019-1065-y

11. Smith BJ, Zhao C, Dubinkina V, Jin X, Moltzau-Anderson J, Pollard KS. Accurate estimation of intraspecific microbial gene content variation in metagenomic data with MIDAS v3 and StrainPGC. bioRxiv. 2024. p. 2024.04.10.588779. doi:10.1101/2024.04.10.588779

12. Page AJ, Cummins CA, Hunt M, Wong VK, Reuter S, Holden MTG, et al. Roary: rapid large-scale prokaryote pan genome analysis. Bioinformatics. 2015;31: 3691–3693. doi:10.1093/bioinformatics/btv421

13. Ding W, Baumdicker F, Neher RA. panX: pan-genome analysis and exploration. Nucleic Acids Res. 2018;46: e5. doi:10.1093/nar/gkx977

14. Song H, Yoo Y, Hwang J, Na Y-C, Kim HS. Faecalibacterium prausnitzii subspecies-level dysbiosis in the human gut microbiome underlying atopic dermatitis. J Allergy Clin Immunol. 2016;137: 852–860. doi:10.1016/j.jaci.2015.08.021

15. Rossi O, Khan MT, Schwarzer M, Hudcovic T, Srutkova D, Duncan SH, et al. Faecalibacterium prausnitzii Strain HTF-F and Its Extracellular Polymeric Matrix Attenuate Clinical Parameters in DSS-Induced Colitis. PLoS One. 2015;10: e0123013. doi:10.1371/journal.pone.0123013

16. Rowe-Magnus DA, Guerout AM, Ploncard P, Dychinco B, Davies J, Mazel D. The evolutionary history of chromosomal super-integrons provides an ancestry for multiresistant integrons. Proc Natl Acad Sci U S A. 2001;98: 652–657. doi:10.1073/pnas.98.2.652

17. Gill SR, Fouts DE, Archer GL, Mongodin EF, Deboy RT, Ravel J, et al. Insights on evolution of virulence and resistance from the complete genome analysis of an early methicillin-resistant Staphylococcus aureus strain and a biofilm-producing methicillin-resistant Staphylococcus epidermidis strain. J Bacteriol. 2005;187: 2426–2438. doi:10.1128/JB.187.7.2426-2438.2005

18. Henke MT, Kenny DJ, Cassilly CD, Vlamakis H, Xavier RJ, Clardy J. Ruminococcus gnavus, a member of the human gut microbiome associated with Crohn’s disease, produces an inflammatory polysaccharide. Proc Natl Acad Sci U S A. 2019;116: 12672–12677. doi:10.1073/pnas.1904099116

19. Zhernakova DV, Wang D, Liu L, Andreu-Sánchez S, Zhang Y, Ruiz-Moreno AJ, et al. Host genetic regulation of human gut microbial structural variation. Nature. 2024;625: 813–821. doi:10.1038/s41586-023-06893-w

20. Fang X, Monk JM, Mih N, Du B, Sastry AV, Kavvas E, et al. Escherichia coli B2 strains prevalent in inflammatory bowel disease patients have distinct metabolic capabilities that enable colonization of intestinal mucosa. BMC Syst Biol. 2018;12: 66. doi:10.1186/s12918-018-0587-5

21. Zhang M, Zheng Y, Sun Z, Cao C, Zhao W, Liu Y, et al. Change in the Gut Microbiome and Immunity by Lacticaseibacillus rhamnosus Probio-M9. Microbiol Spectr. 2023;11: e0360922. doi:10.1128/spectrum.03609-22

22. Martín R, Miquel S, Benevides L, Bridonneau C, Robert V, Hudault S, et al. Functional Characterization of Novel Faecalibacterium prausnitzii Strains Isolated from Healthy Volunteers: A Step Forward in the Use of F. prausnitzii as a Next-Generation Probiotic. Front Microbiol. 2017;8: 1226. doi:10.3389/fmicb.2017.01226

23. Hu W, Gao W, Liu Z, Fang Z, Zhao J, Zhang H, et al. Biodiversity and Physiological Characteristics of Novel Faecalibacterium prausnitzii Strains Isolated from Human Feces. Microorganisms. 2022;10. doi:10.3390/microorganisms10020297

24. Yao S, Zhao Z, Wang W, Liu X. Bifidobacterium Longum: Protection against Inflammatory Bowel Disease. J Immunol Res. 2021;2021: 8030297. doi:10.1155/2021/8030297

25. Breslow NE, Clayton DG. Approximate Inference in Generalized Linear Mixed Models. J Am Stat Assoc. 1993;88: 9–25. doi:10.2307/2290687

26. Chen H, Wang C, Conomos MP, Stilp AM, Li Z, Sofer T, et al. Control for Population Structure and Relatedness for Binary Traits in Genetic Association Studies via Logistic Mixed Models. Am J Hum Genet. 2016;98: 653–666. doi:10.1016/j.ajhg.2016.02.012

27. Zhou W, Nielsen JB, Fritsche LG, Dey R, Gabrielsen ME, Wolford BN, et al. Efficiently controlling for case-control imbalance and sample relatedness in large-scale genetic association studies. Nat Genet. 2018;50: 1335–1341. doi:10.1038/s41588-018-0184-y

28. Loh P-R, Tucker G, Bulik-Sullivan BK, Vilhjálmsson BJ, Finucane HK, Salem RM, et al. Efficient Bayesian mixed-model analysis increases association power in large cohorts. Nat Genet. 2015;47: 284–290. doi:10.1038/ng.3190

29. Loddo I, Romano C. Inflammatory Bowel Disease: Genetics, Epigenetics, and Pathogenesis. Front Immunol. 2015;6: 551. doi:10.3389/fimmu.2015.00551

30. Dahlhamer JM, Zammitti EP, Ward BW, Wheaton AG, Croft JB. Prevalence of Inflammatory Bowel Disease Among Adults Aged ≥18 Years - United States, 2015. MMWR Morb Mortal Wkly Rep. 2016;65: 1166–1169. doi:10.15585/mmwr.mm6542a3

31. Xu F, Carlson SA, Liu Y, Greenlund KJ. Prevalence of Inflammatory Bowel Disease Among Medicare Fee-For-Service Beneficiaries - United States, 2001-2018. MMWR Morb Mortal Wkly Rep. 2021;70: 698–701. doi:10.15585/mmwr.mm7019a2

32. Machiels K, Joossens M, Sabino J, De Preter V, Arijs I, Eeckhaut V, et al. A decrease of the butyrate-producing species Roseburia hominis and Faecalibacterium prausnitzii defines dysbiosis in patients with ulcerative colitis. Gut. 2014;63: 1275–1283. doi:10.1136/gutjnl-2013-304833

33. Wu X, Liu K, Wu Q, Wang M, Chen X, Li Y, et al. Biomarkers of Metabolomics in Inflammatory Bowel Disease and Damp-Heat Syndrome: A Preliminary Study. Evid Based Complement Alternat Med. 2022;2022: 3319646. doi:10.1155/2022/3319646

34. Morgan XC, Tickle TL, Sokol H, Gevers D, Devaney KL, Ward DV, et al. Dysfunction of the intestinal microbiome in inflammatory bowel disease and treatment. Genome Biol. 2012;13: R79. doi:10.1186/gb-2012-13-9-r79

35. Carasso S, Zaatry R, Hajjo H, Kadosh-Kariti D, Ben-Assa N, Naddaf R, et al. Inflammation and bacteriophages affect DNA inversion states and functionality of the gut microbiota. Cell Host Microbe. 2024. doi:10.1016/j.chom.2024.02.003

36. Zhang Y, Bhosle A, Bae S, McIver LJ, Pishchany G, Accorsi EK, et al. Discovery of bioactive microbial gene products in inflammatory bowel disease. Nature. 2022;606: 754–760. doi:10.1038/s41586-022-04648-7

37. Clooney AG, Eckenberger J, Laserna-Mendieta E, Sexton KA, Bernstein MT, Vagianos K, et al. Ranking microbiome variance in inflammatory bowel disease: a large longitudinal intercontinental study. Gut. 2021;70: 499–510. doi:10.1136/gutjnl-2020-321106

38. Dalal SR, Chang EB. The microbial basis of inflammatory bowel diseases. J Clin Invest. 2014;124: 4190–4196. doi:10.1172/JCI72330

39. Sokol H, Seksik P, Furet JP, Firmesse O, Nion-Larmurier I, Beaugerie L, et al. Low counts of Faecalibacterium prausnitzii in colitis microbiota. Inflamm Bowel Dis. 2009;15: 1183–1189. doi:10.1002/ibd.20903

40. Glassner KL, Abraham BP, Quigley EMM. The microbiome and inflammatory bowel disease. J Allergy Clin Immunol. 2020;145: 16–27. doi:10.1016/j.jaci.2019.11.003

41. Vich Vila A, Imhann F, Collij V, Jankipersadsing SA, Gurry T, Mujagic Z, et al. Gut microbiota composition and functional changes in inflammatory bowel disease and irritable bowel syndrome. Sci Transl Med. 2018;10. doi:10.1126/scitranslmed.aap8914

42. Haldar T, Ghosh S. Effect of population stratification on false positive rates of population-based association analyses of quantitative traits: Stratification effects on population-based QTL analyses. Ann Hum Genet. 2012;76: 237–245. doi:10.1111/j.1469-1809.2012.00708.x

43. Hall AB, Yassour M, Sauk J, Garner A, Jiang X, Arthur T, et al. A novel Ruminococcus gnavus clade enriched in inflammatory bowel disease patients. Genome Med. 2017;9: 103. doi:10.1186/s13073-017-0490-5

44. Henke MT, Brown EM, Cassilly CD, Vlamakis H, Xavier RJ, Clardy J. Capsular polysaccharide correlates with immune response to the human gut microbe Ruminococcus gnavus. Proc Natl Acad Sci U S A. 2021;118. doi:10.1073/pnas.2007595118

45. Wood DE, Lu J, Langmead B. Improved metagenomic analysis with Kraken 2. Genome Biol. 2019;20: 257. doi:10.1186/s13059-019-1891-0

46. Lu J, Breitwieser FP, Thielen P, Salzberg SL. Bracken: estimating species abundance in metagenomics data. PeerJ Comput Sci. 2017;3: e104. doi:10.7717/peerj-cs.104

47. Lu J, Rincon N, Wood DE, Breitwieser FP, Pockrandt C, Langmead B, et al. Metagenome analysis using the Kraken software suite. Nat Protoc. 2022;17: 2815–2839. doi:10.1038/s41596-022-00738-y

48. Lavelle A, Nancey S, Reimund J-M, Laharie D, Marteau P, Treton X, et al. Fecal microbiota and bile acids in IBD patients undergoing screening for colorectal cancer. Gut Microbes. 2022;14: 2078620. doi:10.1080/19490976.2022.2078620

49. Nomura K, Ishikawa D, Okahara K, Ito S, Haga K, Takahashi M, et al. Bacteroidetes Species Are Correlated with Disease Activity in Ulcerative Colitis. J Clin Med Res. 2021;10. doi:10.3390/jcm10081749

50. Coyne MJ, Zitomersky NL, McGuire AM, Earl AM, Comstock LE. Evidence of extensive DNA transfer between bacteroidales species within the human gut. MBio. 2014;5: e01305–14. doi:10.1128/mBio.01305-14

51. Kanehisa M, Sato Y, Morishima K. BlastKOALA and GhostKOALA: KEGG Tools for Functional Characterization of Genome and Metagenome Sequences. J Mol Biol. 2016;428: 726–731. doi:10.1016/j.jmb.2015.11.006

52. Lopez-Siles M, Duncan SH, Garcia-Gil LJ, Martinez-Medina M. Faecalibacterium prausnitzii: from microbiology to diagnostics and prognostics. ISME J. 2017;11: 841–852. doi:10.1038/ismej.2016.176

53. Quévrain E, Maubert MA, Michon C, Chain F, Marquant R, Tailhades J, et al. Identification of an anti-inflammatory protein from Faecalibacterium prausnitzii, a commensal bacterium deficient in Crohn’s disease. Gut. 2016;65: 415–425. doi:10.1136/gutjnl-2014-307649

54. Miller KA, Phillips RS, Kilgore PB, Smith GL, Hoover TR. A mannose family phosphotransferase system permease and associated enzymes are required for utilization of fructoselysine and glucoselysine in Salmonella enterica serovar Typhimurium. J Bacteriol. 2015;197: 2831–2839. doi:10.1128/JB.00339-15

55. Bui TPN, Ritari J, Boeren S, de Waard P, Plugge CM, de Vos WM. Production of butyrate from lysine and the Amadori product fructoselysine by a human gut commensal. Nat Commun. 2015;6: 10062. doi:10.1038/ncomms10062

56. Deppe VM, Bongaerts J, O’Connell T, Maurer K-H, Meinhardt F. Enzymatic deglycation of Amadori products in bacteria: mechanisms, occurrence and physiological functions. Appl Microbiol Biotechnol. 2011;90: 399–406. doi:10.1007/s00253-010-3083-4

57. Camargo AP, Roux S, Schulz F, Babinski M, Xu Y, Hu B, et al. Identification of mobile genetic elements with geNomad. Nat Biotechnol. 2023. doi:10.1038/s41587-023-01953-y

58. Singh V, Lee G, Son H, Koh H, Kim ES, Unno T, et al. Butyrate producers, “The Sentinel of Gut”: Their intestinal significance with and beyond butyrate, and prospective use as microbial therapeutics. Front Microbiol. 2022;13: 1103836. doi:10.3389/fmicb.2022.1103836

59. Hodgkinson K, El Abbar F, Dobranowski P, Manoogian J, Butcher J, Figeys D, et al. Butyrate’s role in human health and the current progress towards its clinical application to treat gastrointestinal disease. Clin Nutr. 2023;42: 61–75. doi:10.1016/j.clnu.2022.10.024

60. Yang J, Zaitlen NA, Goddard ME, Visscher PM, Price AL. Advantages and pitfalls in the application of mixed-model association methods. Nat Genet. 2014;46: 100–106. doi:10.1038/ng.2876

61. Pascal V, Pozuelo M, Borruel N, Casellas F, Campos D, Santiago A, et al. A microbial signature for Crohn’s disease. Gut. 2017;66: 813–822. doi:10.1136/gutjnl-2016-313235

62. Zhao C, Shi ZJ, Pollard KS. Pitfalls of genotyping microbial communities with rapidly growing genome collections. Cell Syst. 2023;14: 160–176.e3. doi:10.1016/j.cels.2022.12.007

63. Bolger AM, Lohse M, Usadel B. Trimmomatic: a flexible trimmer for Illumina sequence data. Bioinformatics. 2014;30: 2114–2120. doi:10.1093/bioinformatics/btu170

64. Nurk S, Koren S, Rhie A, Rautiainen M, Bzikadze AV, Mikheenko A, et al. The complete sequence of a human genome. Science. 2022;376: 44–53. doi:10.1126/science.abj6987

65. Breitwieser FP, Pertea M, Zimin AV, Salzberg SL. Human contamination in bacterial genomes has created thousands of spurious proteins. Genome Res. 2019;29: 954–960. doi:10.1101/gr.245373.118

66. Langmead B, Salzberg SL. Fast gapped-read alignment with Bowtie 2. Nat Methods. 2012;9: 357–359. doi:10.1038/nmeth.1923

67. Bushnell B. BBDuk. Jt Genome Inst Available online: https://jgidoegov/data-and-tools/bbtools/bb-tools-userguide/bbduk-guide/(accessed on 25 June 2020). 2020.

68. Babraham Bioinformatics - FastQC A Quality Control tool for High Throughput Sequence Data. [cited 15 May 2024]. Available: https://www.bioinformatics.babraham.ac.uk/projects/fastqc/

69. Efron B. Local False Discovery Rates. 2005. doi:10.1017/cbo9780511761362.006

70. Lee OE, Braun TM. Permutation tests for random effects in linear mixed models. Biometrics. 2012;68: 486–493. doi:10.1111/j.1541-0420.2011.01675.x

71. Zeng P, Zhao Y, Li H, Wang T, Chen F. Permutation-based variance component test in generalized linear mixed model with application to multilocus genetic association study. BMC Med Res Methodol. 2015;15: 37. doi:10.1186/s12874-015-0030-1

72. Benjamini Y, Hochberg Y. Controlling the false discovery rate: A practical and powerful approach to multiple testing. J R Stat Soc. 1995;57: 289–300. doi:10.1111/j.2517-6161.1995.tb02031.x

73. Storey JD. A Direct Approach to False Discovery Rates. J R Stat Soc Series B Stat Methodol. 2002;64: 479–498. doi:10.1111/1467-9868.00346

74. Nelder JA, Wedderburn RWM. Generalized Linear Models. J R Stat Soc Ser A. 1972;135: 370. doi:10.2307/2344614

75. Richardson L, Allen B, Baldi G, Beracochea M, Bileschi ML, Burdett T, et al. MGnify: the microbiome sequence data analysis resource in 2023. Nucleic Acids Res. 2023;51: D753–D759. doi:10.1093/nar/gkac1080

76. Wiame E, Lamosa P, Santos H, Van Schaftingen E. Identification of glucoselysine-6-phosphate deglycase, an enzyme involved in the metabolism of the fructation product glucoselysine. Biochem J. 2005;392: 263–269. doi:10.1042/BJ20051183

77. Camacho C, Coulouris G, Avagyan V, Ma N, Papadopoulos J, Bealer K, et al. BLAST+: architecture and applications. BMC Bioinformatics. 2009;10: 421. doi:10.1186/1471-2105-10-421

78. Seemann T. Prokka: rapid prokaryotic genome annotation. Bioinformatics. 2014;30: 2068–2069. doi:10.1093/bioinformatics/btu153

79. Cantalapiedra CP, Hernández-Plaza A, Letunic I, Bork P, Huerta-Cepas J. eggNOG-mapper v2: Functional Annotation, Orthology Assignments, and Domain Prediction at the Metagenomic Scale. Mol Biol Evol. 2021;38: 5825–5829. doi:10.1093/molbev/msab293

80. Parks DH, Imelfort M, Skennerton CT, Hugenholtz P, Tyson GW. CheckM: assessing the quality of microbial genomes recovered from isolates, single cells, and metagenomes. Genome Res. 2015;25: 1043–1055. doi:10.1101/gr.186072.114

81. Olm MR, Brown CT, Brooks B, Banfield JF. dRep: a tool for fast and accurate genomic comparisons that enables improved genome recovery from metagenomes through de-replication. ISME J. 2017;11: 2864–2868. doi:10.1038/ismej.2017.126

82. Ondov BD, Treangen TJ, Melsted P, Mallonee AB, Bergman NH, Koren S, et al. Mash: fast genome and metagenome distance estimation using MinHash. Genome Biol. 2016;17: 132. doi:10.1186/s13059-016-0997-x

83. Marçais G, Delcher AL, Phillippy AM, Coston R, Salzberg SL, Zimin A. MUMmer4: A fast and versatile genome alignment system. PLoS Comput Biol. 2018;14: e1005944. doi:10.1371/journal.pcbi.1005944

84. Jain C, Rodriguez-R LM, Phillippy AM, Konstantinidis KT, Aluru S. High throughput ANI analysis of 90K prokaryotic genomes reveals clear species boundaries. Nat Commun. 2018;9: 5114. doi:10.1038/s41467-018-07641-9

