## Supplementary Materials for "Improved detection of microbiome-disease associations via population structure-aware generalized linear mixed effects models (microSLAM)"

$$logit(\mu_i) = X_i\alpha + G_i\beta + b_i + \varepsilon_i$$

where $\mu_i = P(y_i = 1 | X_i, G_i, b_i)$ is the probability that the $i$'th individual is a case given the covariates, gene presence/absence vector, and the random effect $b_i$ that is estimated by microSLAM. The random effect $b_i$ is modeled $N(0, \tau\,\psi)$ where $\psi$ the N×N GRM described above, and $\tau$ is the estimated additive genetic variance. The $Var(y_i | b) = \phi\, var(\mu_i)$, in the case of a binary trait the random parameter Φ=1. The parameter $\alpha$ is a $1 \times (1 + p)$ coefficient vector of fixed effects and $\beta$ is a coefficient representing the log odds ratio for the association between the gene's presence and the trait. For a quantitative trait, $y_i$ is a real number and the model is a linear mixed model rather than logistic, so that $\beta$ represents the expected change in the trait for the gene being present versus absent. Everything else is the same.

#### Estimating the coefficients and variance component: fitting microSLAM for each species

We employ the same restricted log-likelihood and average information matrix for estimating the coefficients and variance components as were employed in GMMAT [26] and Saige [27]. For more details on deriving these estimation procedures, refer to those studies and to Clayton and Breslow [25]. We also follow Saige's multi-step process to estimate the random effects and then use these in the logistic (or linear) model presented in the previous section. This helps us in two ways: 1) it reduces computational time significantly as random effects only have to be estimated one time for each species (not once for every gene in every species), and 2) avoiding refitting the random effect for every gene provides a more robust estimate.

Unlike Saige, we do not use PCG, randomized trace estimator, or a low-rank GRM. These are designed to reduce computation and memory costs within the context of human genomes with millions of genetic variants, but these are not major problems for us given the size of the datasets in this study. Also, our GRMs are naturally full-rank. These computational shortcuts could be implemented if needed.

#### Score testing for the GRM: $\tau$ test modeling

We detail microSLAM's $\tau$ test, a new statistical procedure to inform the user whether the species' GRM is significantly related to the trait. This would indicate that a subset of related strains can predict the trait. We consider random effects $b_i \sim N(0, \tau\psi)$, as described above, then compare the models:

$$H_0: Y = X\alpha + \epsilon$$

$$H_a: Y = X\alpha + b + \epsilon$$

After the models have been fit (estimation converges), we have $\hat{\alpha}$; $\hat{b}$; $\hat{\phi}$; and $\hat{\tau}$. We also compute a working vector

$$\hat{Y} = X\hat{\alpha} + \hat{b} + \hat{\epsilon},\ b \sim N(0, \hat{\tau}\psi);\ \hat{\epsilon} \sim N(0, W^{-1})$$

The test statistic for the $\tau$ test can be written as:

$$T = \sum_{i=1}^{N} \hat{b}_i^2 / N$$

This is the sample variance of the estimated random effects $b_i$. This statistic involves the sum of the squared random effect estimates. The null hypothesis is that T=0 (i.e., the random effects do not help to explain variation in the trait). To compute a p-value for T without making assumptions about its distribution, we use a permutation test.

#### Score testing for gene presence/absence: $\beta$ test modeling

After we have fit the model described above for the $\tau$ test, we have estimates of the fixed effect coefficients $\hat{\alpha}$, the random effects $\hat{b}$, and the variance component parameters, $\hat{\phi}$; $\hat{\tau}$. Using these, we construct a score test for each gene with the null hypothesis $H_0 : \beta = 0$. Suppose $G$ is a $N \times 1$ genotype vector (where $N$ is the number of samples). $\hat{\mu}$ are the probabilities of the samples having the trait (e.g., being cases) given the covariates $X$ and the random effects $\hat{b}$: $P(Y = 1 | X, \hat{b})$. Let $\hat{W}$ be a diagonal vector with elements $\hat{\mu}(1 - \hat{\mu})$ and $\tilde{G} = G - X(X^t\hat{W}X)^{-1}X^t\hat{W}G$ is the covariate-adjusted genotype vector. With $\hat{\Sigma} = \hat{W}^{-1} + \hat{\tau}\psi$ and

$P = \hat{\Sigma}^{-1} - \hat{\Sigma}^{-1}X(X^t\hat{\Sigma}^{-1}X)^{-1}X^t\hat{\Sigma}^{-1}$ and a working vector $\hat{Y} = X\hat{\alpha} + \hat{b}_i + g'(\hat{\mu})(y - \hat{\mu})$, the score test statistics, assuming $\hat{P}\tilde{G} = \hat{P}G$ is:

$$T = G^t(Y - \hat{\mu}) = G^t\hat{P}\hat{Y} = \tilde{G}^t\hat{P}Y = \tilde{G}^{\hat{t}}(Y - \hat{\mu})$$

The variance of T is:

$$Var(T) = \tilde{G}W\tilde{G}$$

We estimate this directly for each gene $G$. As shown in [27] this is approximately equivalent to $\tilde{G}\hat{P}\tilde{G}$ but much faster to compute, plus the approximation is conservative.

The effect size $\hat{\beta}$ is the natural log of the odds ratio. We can estimate this using the variance component estimate under the null hypothesis.

$$\hat{\beta} = (\tilde{G}W\tilde{G})^{-1}(\tilde{G}\hat{P}\hat{Y}) = T/var(T)$$

The standard error of $\hat{\beta}$ is $SE(\hat{\beta}) = \left|\hat{\beta}/z\right|$ where $z$ is the z-score corresponding to the p-value divided by 2.

#### Simulations

For the β test simulation 3, we sought to generate gene presence/absence matrices, trait values, and age values without using any real sample data. First, we simulated a trait from the binomial distribution with a success probability of 0.5. We assume that there are two strains, one that is correlated with the trait ($odds\ ratio = 2.22$) and one uncorrelated with it. For the correlated strain, we simulated 300 genes at a presence level of 0.5 and an odds ratio of 4.0 for the gene being present given a person has the strain. We simulated one-third of the samples having the uncorrelated strain and modeled this with 250 genes with low binomial presence rates ($p = 0.3$) and an odds ratio of 4.0 for the gene being present given a person has the strain. Then we modeled 300 genes that were “core” across all samples; these were drawn from a binomial with a success probability of 0.8. We additionally simulated 150 genes that were non-strain associated “accessory” genes; these were drawn from a binomial success probability of 0.2. Last, we simulated at least one gene ($G_y$) that is even more highly correlated with the trait than is the correlated strain ($odds\ ratio = 2.44$). The more genes in $G_y$ (we investigated 1, 2, or 3 genes) and the stronger the relationship between $G_y$ and the trait, the higher the parameter $\tau$ will be. The resulting gene presence/absence matrices naturally have a range of different values of $\tau$. Age was randomly generated with parameters similar to the IBD data: $ceiling(rnorm(N, mean = 45, sd = 15))$. We repeated Simulation 3 with the number of samples varying from 60 to 250.

### Supplemental Figures

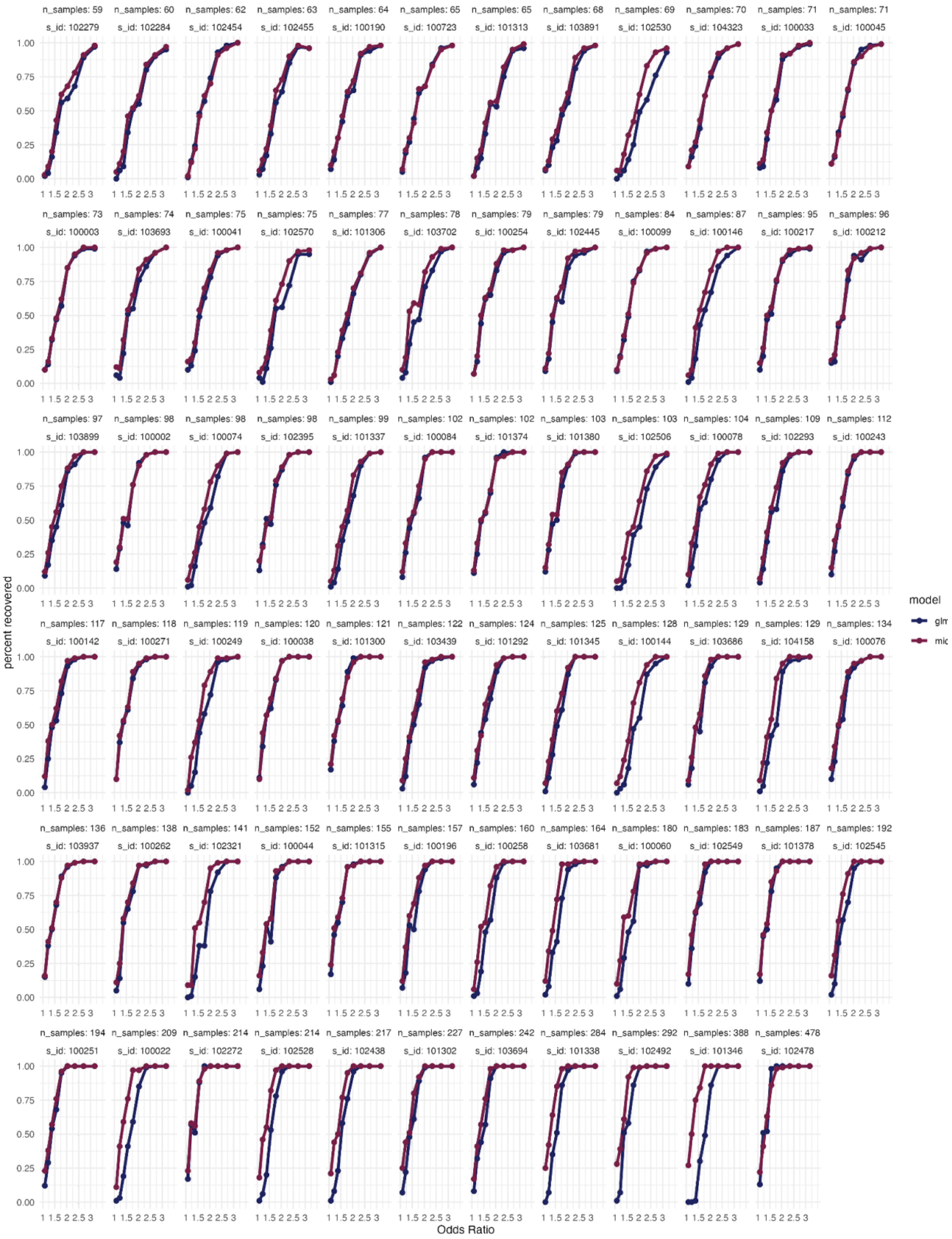
n_samples: 59 s_id: 102279
n_samples: 60 s_id: 102284
n_samples: 62 s_id: 102454
n_samples: 63 s_id: 102455
n_samples: 64 s_id: 100190
n_samples: 65 s_id: 100723
n_samples: 65 s_id: 101313
n_samples: 68 s_id: 103891
n_samples: 69 s_id: 102530
n_samples: 70 s_id: 104323
n_samples: 71 s_id: 100033
n_samples: 71 s_id: 100045
n_samples: 73 s_id: 100003
n_samples: 74 s_id: 103693
n_samples: 75 s_id: 100041
n_samples: 75 s_id: 102570
n_samples: 77 s_id: 101306
n_samples: 78 s_id: 103702
n_samples: 79 s_id: 100254
n_samples: 79 s_id: 102445
n_samples: 84 s_id: 100099
n_samples: 87 s_id: 100146
n_samples: 95 s_id: 100217
n_samples: 96 s_id: 100212
n_samples: 97 s_id: 103899
n_samples: 98 s_id: 100002
n_samples: 98 s_id: 100074
n_samples: 98 s_id: 102395
n_samples: 99 s_id: 101337
n_samples: 102 s_id: 100084
n_samples: 102 s_id: 101374
n_samples: 103 s_id: 101380
n_samples: 103 s_id: 102506
n_samples: 104 s_id: 100078
n_samples: 109 s_id: 102293
n_samples: 112 s_id: 100243
n_samples: 117 s_id: 100142
n_samples: 118 s_id: 100271
n_samples: 119 s_id: 100249
n_samples: 120 s_id: 100038
n_samples: 121 s_id: 101300
n_samples: 122 s_id: 103439
n_samples: 124 s_id: 101292
n_samples: 125 s_id: 101345
n_samples: 128 s_id: 100144
n_samples: 129 s_id: 103686
n_samples: 129 s_id: 104158
n_samples: 134 s_id: 100076
n_samples: 136 s_id: 103937
n_samples: 138 s_id: 100262
n_samples: 141 s_id: 102321
n_samples: 152 s_id: 100044
n_samples: 155 s_id: 101315
n_samples: 157 s_id: 100196
n_samples: 160 s_id: 100258
n_samples: 164 s_id: 103681
n_samples: 180 s_id: 100060
n_samples: 183 s_id: 102549
n_samples: 187 s_id: 101378
n_samples: 192 s_id: 102545
n_samples: 194 s_id: 100251
n_samples: 209 s_id: 100022
n_samples: 214 s_id: 102272
n_samples: 214 s_id: 102528
n_samples: 217 s_id: 102438
n_samples: 227 s_id: 101302
n_samples: 242 s_id: 103694
n_samples: 284 s_id: 101338
n_samples: 292 s_id: 102492
n_samples: 388 s_id: 101346
n_samples: 478 s_id: 102478
0.00 0.25 0.50 0.75 1.00
1 1.5 2 2.5 3
percent recovered
Odds Ratio
model

**Supplemental Figure S1 - GLM and microSLAM β test power evaluations for 71 simulated species.** These plots show estimated power of β tests using data from simulation 2 in which a gene presence/absence matrix and binary trait were simulated based on the observed GRMs from the 71 species in the IBD compendium using a range of different effect sizes (odds ratios, horizontal axes). There is one panel per GRM (labeled with species ID), and panels are ordered from lowest to highest sample size.Power was computed as the proportion of positive genes discovered at an empirical localFDR of 0.05 for both microSLAM (*red*) and glm (*blue*). As the number of samples increases there tends to be a larger difference between the glm and the microSLAM models.

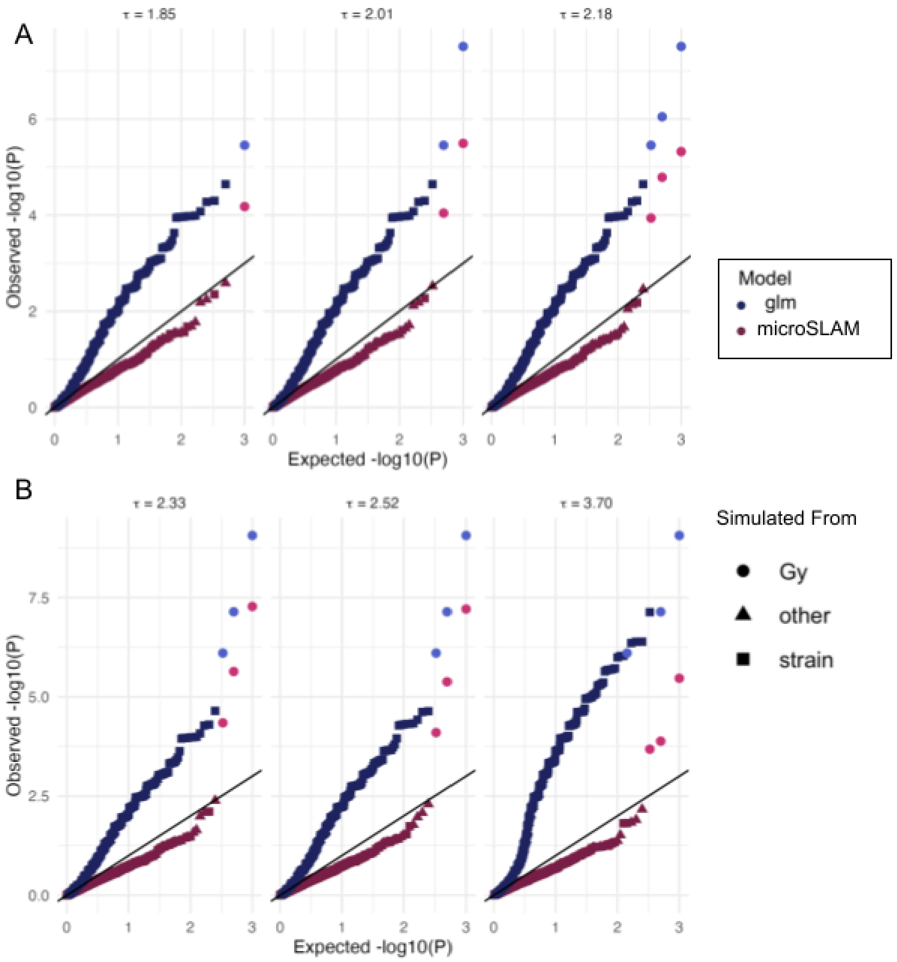


**Supplemental Figure S2 - P-values from microSLAM's β test are somewhat conservative, while GLM's are inflated.** Q-Qplots of β Test results for a simulation with positive genes ($G_y$; *pink circles*: microSLAM, *light blue circles*: glm) plus negative genes that are linked to a strain (strain; squares) or randomly generated (other; triangles) (β test simulation 3, **Supplemental Text).** A) Compared to glm (blue), microSLAM (red) better distinguishes the positive genes $G_y$ from those simulated from the strain. The number of positive genes was one (*left*), two (*middle*), or three (*right*). The value of $\tau$ increases with each additional gene $G_y$. B) As the relationship between the strain and *y* is increased (left to right), the value of $\tau$ increases, and the rate of inflation increases for glm. Across different values of $\tau$, microSLAM remains slightly conservative and continues to rank the positive genes $G_y$ highest, indicating high specificity.

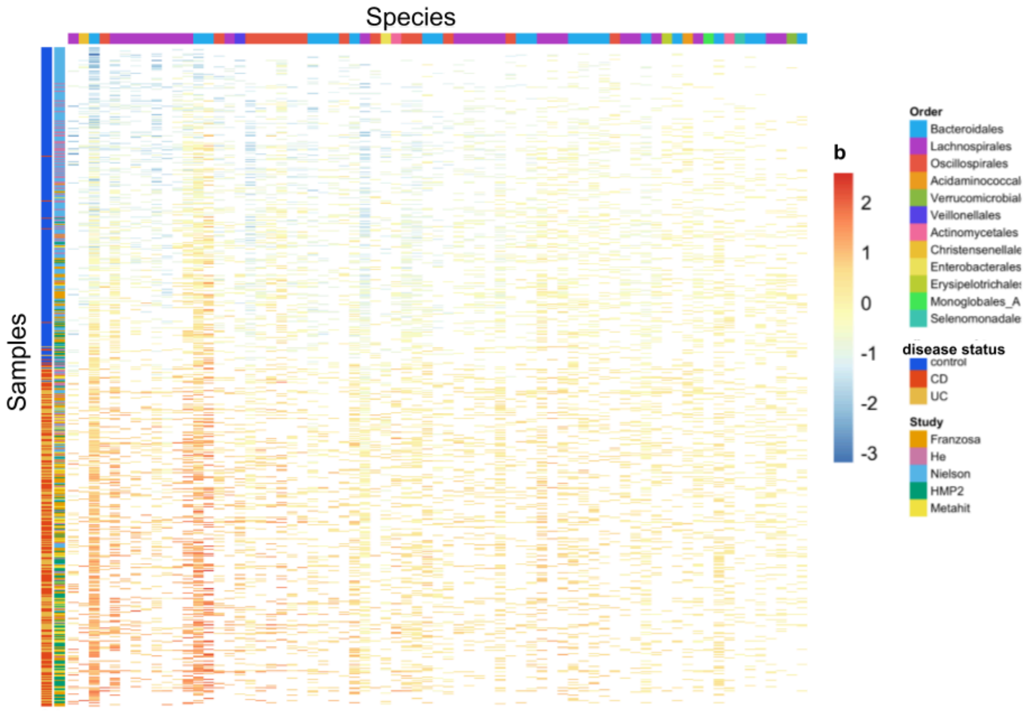


**Supplemental Figure S3 - Random effect (b) values across studies and species**. This heatmap shows microSLAM estimates of the random effect parameters (b) for each of 71 species across all samples where it was detected in the IBD compendium. The red-to-blue color scale denotes the association between strains and IBD (binary case/control status). Red: strains positively associated with IBD; Blue: strains negatively associated with IBD. The study and IBD subtype of each sample are shown on the left. IBD subtypes: Blue=control, red=Crohn's disease (CD), yellow=Ulcerative colitis (UC). CD and UC were combined as cases in the microSLAM modeling. Studies: Franzosa (NCBI BioProject PRJNA400072; orange), He (PRJNA398089; pink), Nielsen (PRJEB15371; blue), HMP2 (PRJEB5224; green), MetaHIT (PRJEB1220; yellow). Species are ordered by the standard deviation of *b* (left=highest standard deviation), where higher standard deviation indicates greater strain diversity that is associated with case/control status. The samples in each column are ordered by lowest to highest average b value. A few studies (e.g., Nielsen) have more controls than others, but there is no systematic relationship between study and population structure. CD and UC tend to have similar distributions of *b* values (i.e., red and yellow are mixed on the left side bar). While we cannot rule out confounders that were unmeasured in the publicly available data that we could access, these patterns suggest that our findings are not obviously biased by differences in study population (e.g., diet, medical care, geography, type of IBD) that could confound measured associations between case/control status and microbiome strains and genes.

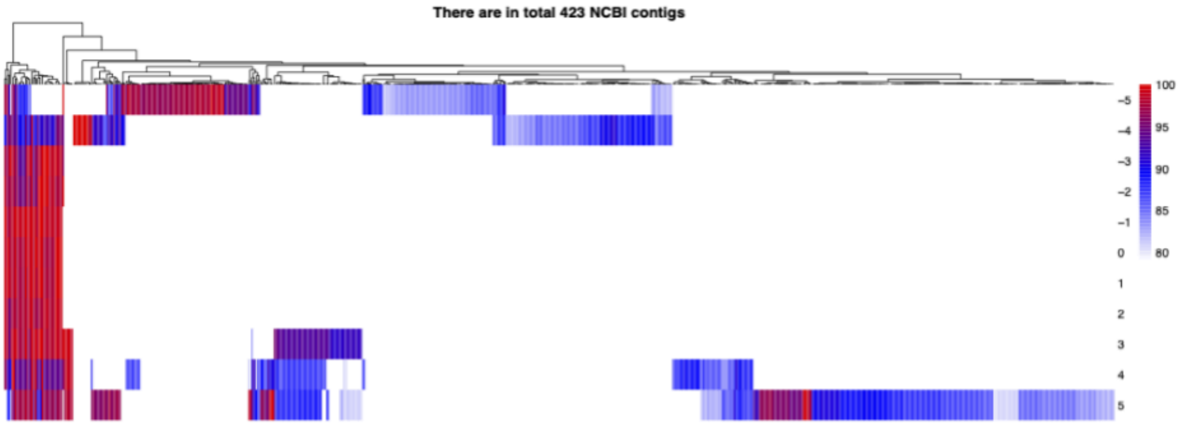


**Supplemental Figure S4 - *F. prausnitzii PTS* operon evolves as a unit across diverse genomes.** This operon comprises seven genes (occasionally eight genes) that were consistently present or absent together across 53% (49/85) of *F. prausnitzii* genomes from NCBI. The order and orientation of genes in the operon is conserved. This heatmap shows the genes (rows; position 0 is *gfrD,* which was significant after localFDR adjustment of microSLAM β test p-values). The other genes were significant before localFDR adjustment and are indexed relative to *gfrD* in the heatmap. Columns represent 423 contigs from 85 *F. prausnitzii* high-quality NCBI genomes. The color of the heatmap shows the blastn sequence similarity of the gene sequence in the contig compared to the sequence in the *F. prausnitzii* reference genome used in our microSLAM analysis (red=highest similarity, white=no significant match). The seven genes in the operon (middle rows of the heatmap) have high sequence similarity when they are present and are present together (red on left), whereas flanking genes are more variably present and have lower sequence identity (blue in top and bottom rows).

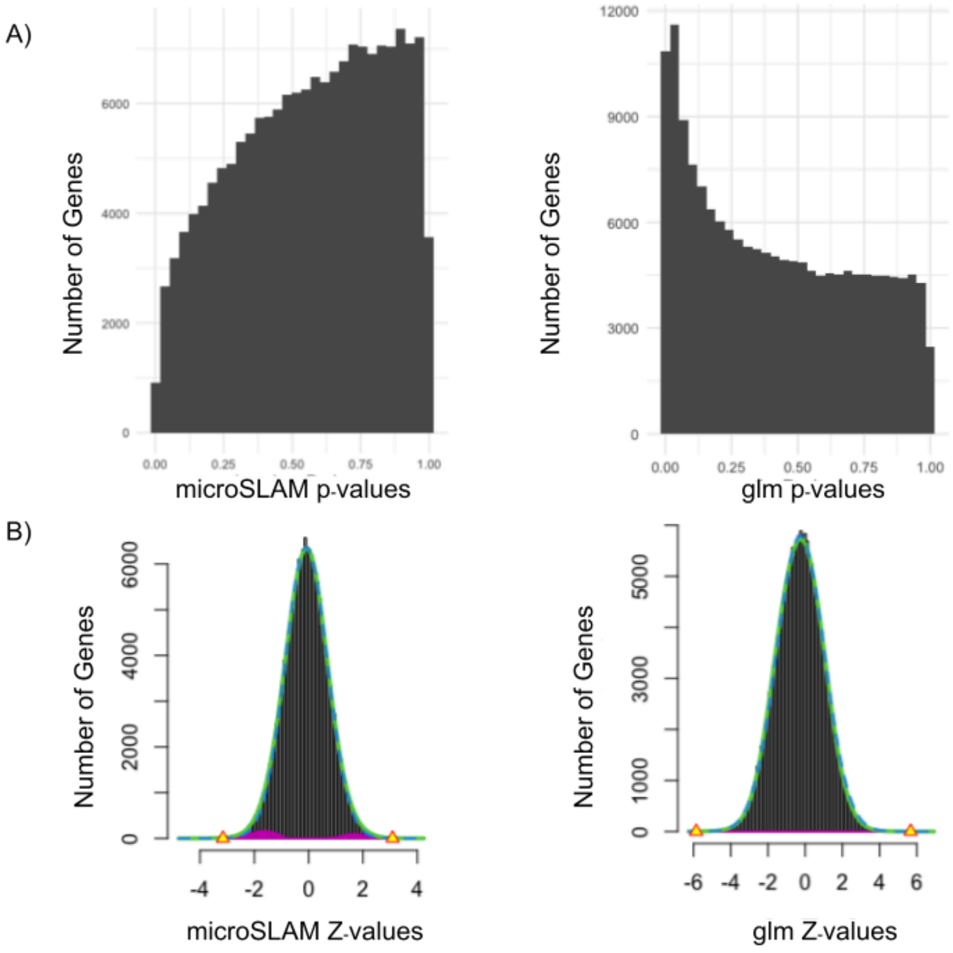


**Supplemental Figure S5 - LocalFDR p-values and Z-values**. A) Histogram of p-values for microSLAM's β test (left) and glm (right). B) Output from localFDR showing the distribution of the null z-values (green) versus the distribution of the z-values that do not follow the null (pink). Yellow triangles denote the z-value thresholds corresponding to a localFDR of 0.2.

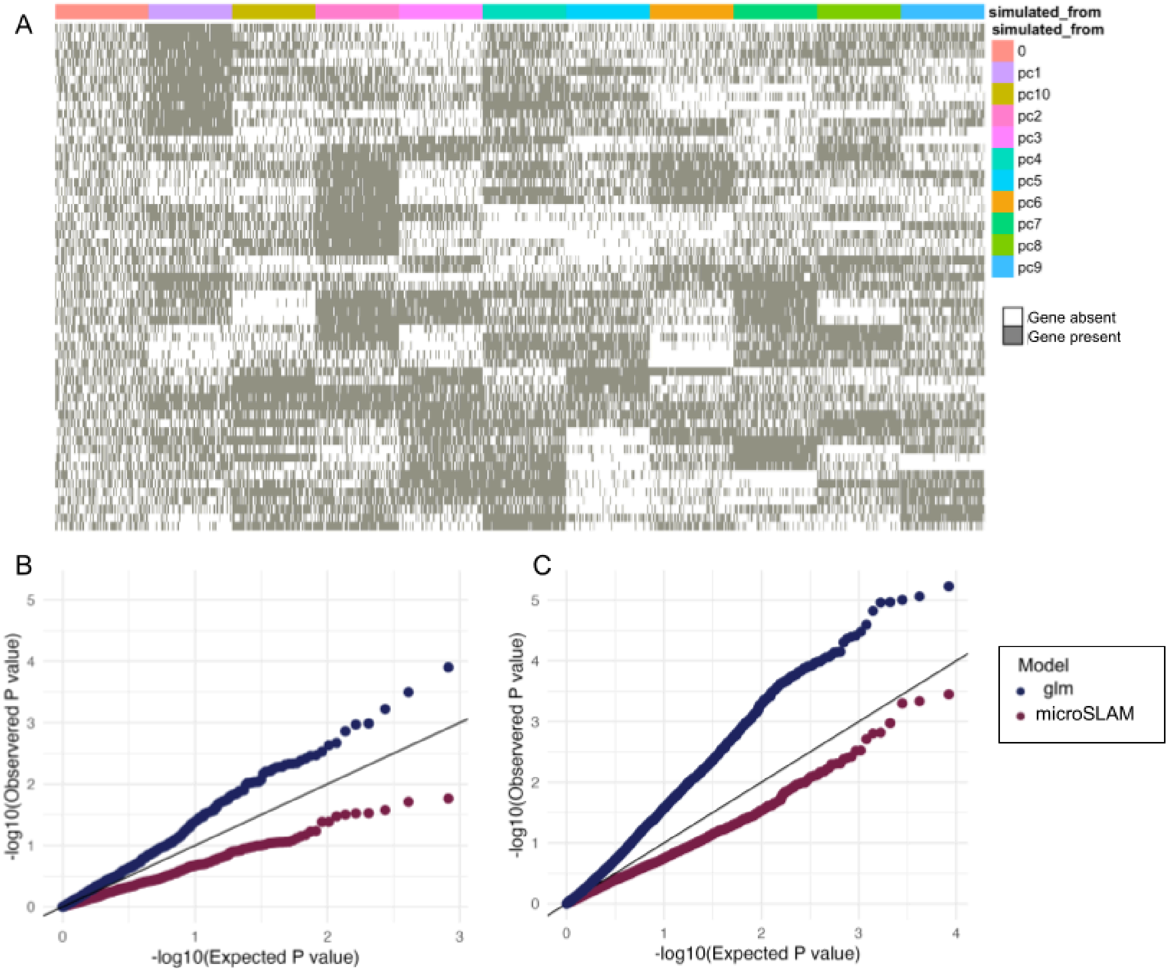


**Supplemental Figure S6 - Example of data and results from microSLAM β test Simulation 1**. A) Simulated gene presence/absence matrix based on the GRM of *Bacteroides thetaiotaomicron* plotted as a heatmap (grey=gene present in a given sample, white=gene absent). Genes are in columns and are labeled according to how they were simulated (0=random, pc1-10=using one of the first 10 principal components of the observed GRM for *B. thetaiotaomicron*. This presence/absence matrix has a some population structure (estimated $\tau$ = 2.30), but no genes were simulated to be associated directly associated with the trait which is defined by the first two PCs. B) Q-Qplot of p-values from all genes not from PC1 or 2 from microSLAM's **β** test (red) and glm (blue) applied to the simulated gene presence/absence matrix in (A). There is a much higher error rate for the glm model. On the other hand, microSLAM is overly conservative (i.e., underpowered). C) Q-Q plot for microSLAM's $\boldsymbol{\beta}$ test (red) and glm (blue) applied to the observed *B. thetaiotaomicron* gene presence/absence matrix from the IBD compendium. The trends are very similar to those in the simulation.

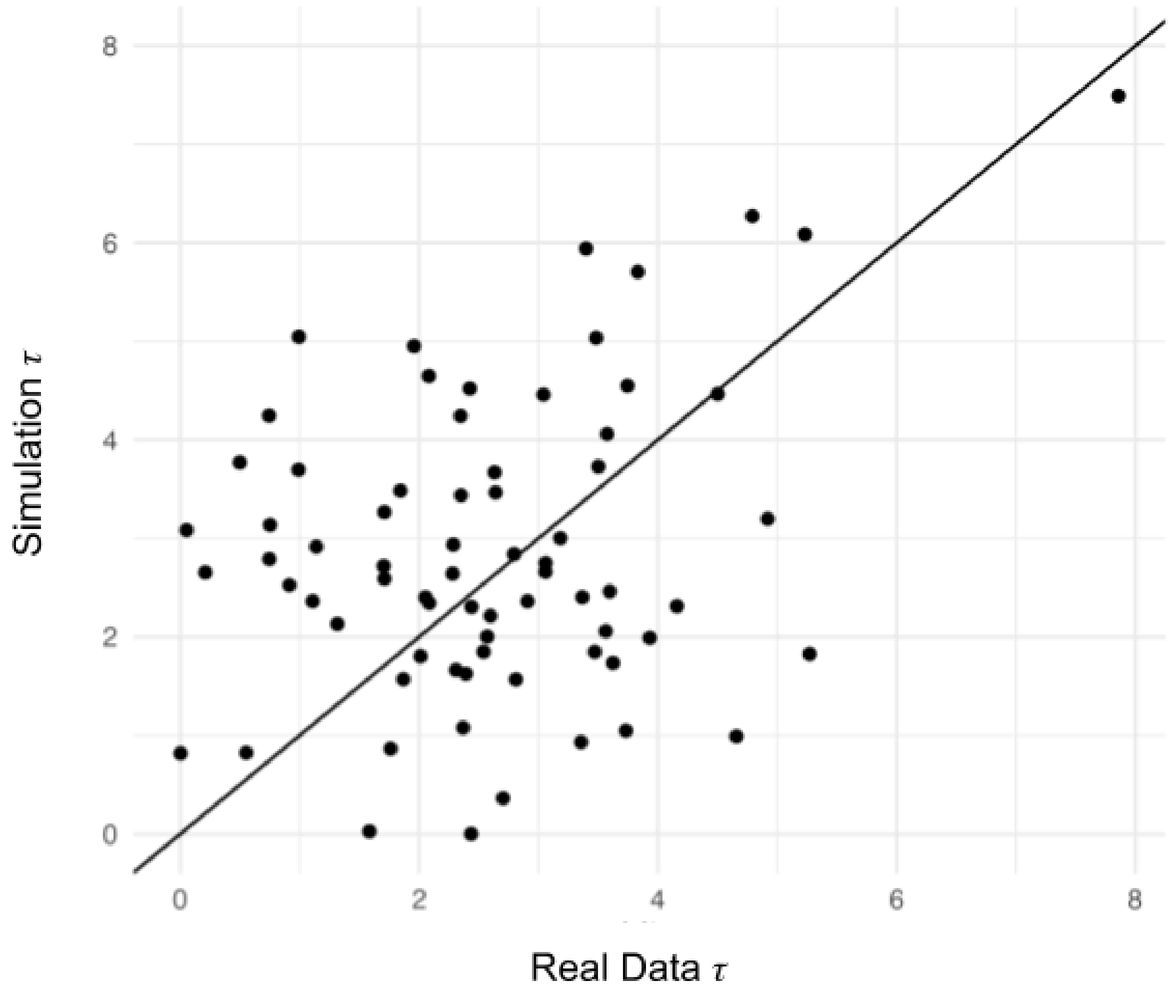


**Supplemental Figure S7 - β Test simulation τ values versus observed τ values in IBD compendium** In the β Test Simulation 1 and 2 set up, we generated gene presence/absence matrices using the observed GRMs for the 71 species in the IBD compendium. Our objective was to generate simulated data that was similar to but not identical to the observed data (**Methods**). This scatter plot shows the $\tau$ values estimated by microSLAM on the simulated data (*y axis)* compared to the corresponding $\tau$ values estimated from the real data in the IBD compendium (*x axis*). The n $\tau$ from the simulation cover a similar range of values as those from the real data while not being highly correlated.
